# Disrupted glial-mediated synaptic refinement in Fragile X syndrome

**DOI:** 10.64898/2026.04.10.717772

**Authors:** Lindsey Starr, Melissa Lee, Amy Vo, Maia Weisenhaus, Lucas Cheadle, Archana Yadav, Fahad Paryani, Mimi Shirasu-Hiza, Vilas Menon, Carol Mason

## Abstract

Fragile X syndrome (FXS), the most common inherited cause of intellectual disability and autism, results from the loss of the RNA-binding protein fragile X mental retardation protein (FMRP). FMRP is a translational regulator and is highly expressed in glial cells, where its role in neural circuit development remains poorly defined. Here, it was observed that *Fmr1* knockout mice exhibit reduced synapse size and accelerated eye-specific segregation. To examine which cell-types participate in this process, a multi-omic framework was applied to FXS model mice at postnatal day 7, a critical window for synaptic remodeling in the retinogeniculate pathway, an established model system utilized to study synaptic pruning. Single-cell transcriptomics revealed coordinated alterations in microglia, astrocytes, and neurons in genes linked to synaptic pruning. Computational modeling further demonstrated enhanced astrocyte-to-microglia signaling, particularly through Ephrin A (EphA)- and semaphorin-mediated pathways, while lipidomic profiling revealed reductions in EphA-associated lipid species required for lipid raft stability and receptor localization. Consistent with these observations, a glial engulfment assay indicated that FXS microglia and astroglia over-engulf synaptic material in the lateral geniculate nucleus, supporting the transcriptomic profile. Together, these findings identify impaired glial-driven synaptic refinement as an early mechanistic feature of FXS pathogenesis, highlighting the genes involved in this process as potential therapeutic targets during circuit development.

## INTRODUCTION

Fragile X syndrome (FXS) is a neurodevelopmental disorder that represents the most common monogenic cause of both autism and intellectual disability. The vast majority of FXS cases are caused by a trinucleotide repeat expansion of CGG within the 5’ untranslated region (UTR) of the fragile X mental retardation 1 (*FMR1*) gene,^1^ resulting in *FMR1* gene silencing and loss of the protein product, fragile X mental retardation protein (FMRP). FMRP is a RNA-binding protein that acts canonically as a mRNA translocator, mRNA stabilizer, and negative translational regulator, repressing up to 4% of total fetal human mRNAs.^2–5^ Emerging evidence suggests there are additional roles of FMRP, including regulation of genome stability,^6,7^ pre-mRNA alternative splicing,^8^ ion channel gating,^3,9^ and modulating cell differentiation kinetics.^10^ In some cases, FMRP can also act as a translational activator.^1,12^ Collectively, these findings highlight FMRP as a central regulator of neural development and function, with potential implications not only for neurons, but also for glial populations in the brain.

In mice, FXS has been modeled using the *Fmr1* knockout (KO) mouse, which lacks expression of FMRP and exhibits many of the synaptic, behavioral, and molecular abnormalities observed in human patients.^13^ The mouse retinogeniculate circuit is also a well-characterized model of activity-dependent synapse maturation that provides a powerful system for studying synaptic refinement. During normal retinogeniculate development, retinal ganglion cell (RGC) axons extend from each eye to the dorsal lateral geniculate nucleus (dLGN) of the brain thalamus, where the ipsilateral (same side) and contralateral (opposite side) axons initially form overlapping projections. The excess synapses are then selectively eliminated via activity-dependent pruning. This is completed by postnatal day 12 (P12), such that each dLGN neuron only receives input from a single eye, either ipsilateral or contralateral.^14–17^ Thus, the retinogeniculate system in *Fmr1* KO mice provides a powerful *in vivo* model to study whether and how the loss of FMRP impairs synaptic pruning and to identify how specific cell-types, especially glia, contribute to abnormal circuit development in FXS.

Glia, the non-neuronal cells of the brain, have a critical role in developmental synapse elimination. Specifically, astrocytes, microglia, and oligodendrocyte precursor cells (OPCs) actively engulf weaker synapses during periods of neural circuit refinement.^18–22^ Although FXS research has traditionally centered on neuronal dysfunction, both *Fmr1* and FMRP are highly expressed in glial cell-types^23–25^; thus, glia may have an important role in disease formation. Indeed, recent studies using cell-type-specific deletion and re-expression of *Fmr1* in mice suggest that astrocytes contribute to multiple FXS-related phenotypes, including abnormal dendritic spine structure, neuronal hyperexcitability, deficits in social novelty preference, and impaired memory.^26,27^ Moreover, studies in *Drosophila* models of FXS indicate that ensheathing glia exhibit impaired synaptic pruning.^28^ These findings indicate that glia are active contributors to the synaptic and behavioral abnormalities associated with FXS, yet their role in disease pathogenesis remains largely unexplored.

Despite increasing evidence that glia contribute to synapse elimination and are affected by FMRP loss, it remains unclear whether glial-mediated synaptic pruning is disrupted in FXS. Moreover, the specific molecular mechanisms by which FMRP regulates glial function during synaptic refinement have not been clearly defined. To address this gap in knowledge, this study employs a multidisciplinary approach that combines high-resolution imaging, single-cell transcriptomics, computational modeling of intercellular communication, lipidomic profiling, and quantitative analysis of glial engulfment. Here, it was found that *Fmr1* KO mice exhibit reduced synapse size and altered segregation in the development dLGN at P7. This was accompanied by the differential expression of genes associated with synaptic refinement across neurons and glia. Computational modeling of cell-type communication indicated increased astrocyte-to-microglia signaling in the *Fmr1* KO dLGN, driven by multiple lipid pathways. Preliminary lipidomic profiling detected shifts in brain and serum lipid composition, particularly in Ephrin A (EphA)-associated lipid species, in *Fmr1* KO mice, which may be restored via Lovastatin treatment. Finally, both astrocytes and microglia displayed significantly enhanced engulfment of synapses in the absence of FMRP. Together, these findings establish a previously unrecognized role for FMRP in coordinating glial-driven synaptic remodeling through transcriptional and intercellular mechanisms, highlighting glial dysfunction as a key contributor to developmental abnormalities in FXS.

## MATERIALS AND METHODS

### Animals

Mice were maintained in barrier facilities at Columbia University Irving Medical Center and Columbia University Jerome L. Greene Science Center on a 12-hr light-dark cycle. All procedures were carried out in compliance with protocols approved by Columbia University’s Institutional Animal Care and Use Committee. Aldh1l1-eGFP mice were generated by GENSAT^29^ and were provided as a gift from Dr. Ben Barres (Stanford University). *Fmr1* KO mice (B6.129P2-*Fmr1*tm1Cgr/J; strain 003025) and WT mice (C57BL/6J; strain 000664) were obtained from The Jackson Laboratory. As Fragile X syndrome primarily affects males, all experiments were performed exclusively on male littermates. To produce *Fmr1* KO or *Fmr1* WT littermate males, *Fmr1* KO females were crossed to either WT or Aldh1l1-eGFP+/+ males. All mice were maintained on a C57BL/6J background. All experimental procedures and analyses were performed blind to genotype unless otherwise noted.

### Synapse labeling, imaging, and analysis

P7 *Fmr1* KO and WT mice were anesthetized by intraperitoneal injection of ketamine/xylazine, transcardially perfused with PBS, and heads fixed in 4% PFA overnight. Brains were dissected out, cryoprotected serially in 10% and 30% sucrose-PBS solutions, and cryosectioned at 12 *μ*M thickness. Synaptic immunohistochemistry, imaging, and analysis were performed as described.^30^ Briefly, sections were blocked in 10% NGS-PBS solution, followed by overnight incubation in primary antibody (VGlut2, Homer) and a 2-hr secondary antibody incubation. Three sections containing medial dLGN 60 per animal were imaged for further analysis. Two field-of-view Z-stacks targeting the putative ipsilateral-contralateral RGC overlap zone in the dLGN were taken per section on a Nikon A1R Confocal microscope, using the 63X oil objective (NA: 1.4, CFI60 Plan Apochromat *λ*). Images were processed in FIJI and analyzed using the ImageJ Puncta Analyzer plug-in (written by Bary Wark).

### RGC segregation assay

P6 *Fmr1* KO and WT mice were anesthetized by hypothermia, and a micro-feather ophthalmic scalpel (PFM Medical, 200300745) was used to make an incision along the developing eye-fold. Fine-point forceps (Fisher Scientific, 12-000-126) were used to gentry extrude the eye, and 1-2 *μ*l of cholera toxin *β*-subunit (CTB) conjugated to Alexa488 and Alexa594 (Invitrogen, 1 mg/ml with 1% DMSO) was injected intravitreally into the left and right eyes via glass micropipette, and the mouse were allowed to recover. At P7 mice were transcardially perfused with a 4% paraformaldehyde (PFA) solution and further fixed in 4% PFA overnight. Following perfusion, brains were dissected, embedded in 3% agarose, and coronal vibratome sections of 100 *μ*M thickness were taken throughout the dLGN. The three medial-most dLGN sections were imaged on a Zeiss AxioImager M2 Microscope with Apotome, AxioCam MRm camera, using a 10x objective and Neurolucida software (v11, MBF Biosciences, Williston, VT, USA, RRID:SCR_001775). Images were acquired in MBF .tiff format and were processed into 8-bit image files in Fiji (ImageJ, NIH, RRID: SCR_002285). Animals with visible RGC lesioning in the dLGN were excluded from analysis. Threshold-independent segregation analysis was performed as previously described^31,32^ using Metamorph software. Statistical comparison was performed using grouped analysis of multiple t-tests, with Holm-Sidak correction for multiple comparisons.

### Single-nucleus RNA sequencing, data processing, and clustering

P7 *Fmr1* KO and WT mice were anesthetized by intraperitoneal injection of ketamine/xylazine, quickly decapitated, and their brains dissected. 300 μM vibratome sectioning throughout the brain was performed in ice-cold PBS, and dLGN volumes were quickly micro dissected using a micro-knife with a needle blade (Fine Science Tools, 10318-14) and flash-frozen in liquid nitrogen. Single nuclei were dissociated and barcoded using the inDrops platform.^33^ Libraries were created as previously described,^33–36^ indexed, pooled, and sequenced on a NextSeq 500 (Illumina).

Sequenced reads were processed using a previously published pipeline.^37^ Briefly, Bowtie 1.1.1 was used to build a custom transcriptome from Ensembl GRCm38 genome and GRCm38.84 annotation, after filtering the annotation gtf file (gencode.v17.annotation.gtf filtered for feature_type = “gene,” gene_type = “protein_coding” and gene_status = “KNOWN”), using otherwise de- fault parameters. Quality control of reads were performed against this transcriptome, along with read mapping. Sequence reads were linked back to individual captured molecules using unique molecular identifiers (UMIs).

Data from all nuclei with < 90% mitochondrial content and UMI counts between 500 and 150,000 were consolidated into a single dataset and clustered using the Seurat3 R package.^38^ Data were log normalized and scaled to 10,000 transcripts per cell, and variable genes were identified with the following parameters: x.low.cutoff = 0.0125, x.high.cutoff = 3, y.cutoff = 0.5. Analysis was limited to the top 30 principal components (PCs), and clustering resolution was set to 0.5. Cell type clusters were identified by examining known marker gene expression, with *Snap25* with *Slc17a6*, *Snap25* with *Gad1, Olig1, Aqp4, Cx3cr1, Cldn5*, and *Vtn* used to identify excitatory neurons, inhibitory neurons, oligodendrocytes, astrocytes, microglia, endothelial cells, and pericytes, respectively.

### Pseudobulking and differential expression analysis

To enable differential expression at the cell-type level, single-nucleus RNA sequencing data were aggregated across nuclei from the same cell-type and individual mouse, using a pseuodbulking approach. For each cell-type including microglia, astrocytes, excitatory neurons and inhibitory neurons, gene-level UMI counts from all nuclei belonging to a given mouse were summed to generate one count vector per gene, per donor. This pseuodbulking strategy reduces technical noise and enables statistical testing across biological replicates, while still preserving cell-type specificity.

Cell-types with insufficient representation per sample were excluded from downstream analysis to ensure statistical reliability.^39^ Comparisons between *Fmr1* KO and WT mice were conducted separately for each cell-type using established methods for normalization and dispersion estimation. Genes were considered significantly differentially expressed if they met thresholds for statistical significance (adjusted p-value < 0.05) and effect size (absolute log, fold change ≥ 0.25), following multiple testing correction.

### Gene Ontology Analysis

Gene ontology (GO) enrichment analysis was performed using the topGO R^40^ package to identify biological processes overrepresented among differentially expressed genes in the *Fmr1* KO dLGN. GO analysis was conducted separately for each cell-type including microglia, astrocytes, excitatory neurons and inhibitory neurons, using the sets of genes identified as DEGs from the DESeq2 analysis. The background gene set for each cell-type included all genes detected in the corresponding pseudobulk gene expression data. Enrichment was assessed via the topGO “weight01” algorithm, which reduces the influence of broad, generic terms to highlight more specific biological processes. Fisher’s exact test was utilized to determine statistical significance of enrichment and GO terms with a p-value < 0.05 were considered significantly enriched.

### Cell-Cell communication analysis and total interaction potential (TIP) calculation

To investigate changes in intercellular signaling between *Fmr1* KO and WT mice, cell-cell communication analysis was performed using the CellChat R package.^41^ Pseudobulked, log-normalized gene expression matrices were generated separately for microglia, astrocytes, excitatory neurons, and inhibitory neurons from WT and *Fmr1* KO mice. Separate CellChat objects were created for each genotype, and the curated mouse-ligand-receptor interaction database was used to infer signaling interactions. Overexpressed ligands and receptors were identified, and intercellular communication probabilities were estimated using a truncated mean approach. Communication networks were filtered to exclude low confidence interactions and were projected onto a protein-protein interaction network to enhance biological interpretability.

For each genotype, CellChat computed global interaction matrices, pathway-specific communication strength, and source-target hierarchies. The total interaction potential (TIP) was calculated as the sum of all predicted ligand-receptor interactions between a given source and target cell-type, integrating both the number and strength of individual signaling events. This metric provides a quantitative estimate of the overall communication capacity between pairs. Comparative analysis between *Fmr1* KO and WT mice was performed by computing TIP values across all source-target combinations and signaling pathways. Differences in TIP scores were used to identify which cell-types had altered communication in *Fmr1* KO mice and altered signaling pathways were ranked based on the magnitude of change in their overall interaction strength compared to WT controls.

### Lipidomic analysis

Lipidomic profiling was conducted on brain and serum samples from mice treated with pharmaceutical grade Lovastatin across four experimental conditions: WT untreated, *Fmr1* KO untreated, WT treated, *Fmr1* KO treated. Lovastatin is an FDA-approved, widely used cholesterol-lowering drug that acts as a HMG-CoA reductase inhibitor and that has been shown to ameliorate FXS-associated phenotypes when given to *Fmr1* KO mice. The FXS symptoms include excess protein synthesis in the brain (30-100 mg/kg acute treatment via IP injection^42^), audiogenic seizure susceptibility (100 mg/kg via IP injection)^43^, and associative learning defects (100 mg/kg via rodent chow)^44^. Mice received the drug daily intraperitoneally at a dose of 50 mg/kg at 0.1, 3, 24 hours, from postnatal day 4 to postnatal day 6. All mice were fed a standard chow diet throughout the study. Mice were sacrificed at postnatal day 8, and brain and serum samples were harvested immediately, flash-frozen in liquid nitrogen, and stored at −80 °C until analysis. For each condition, six biological replicates per tissue type (brain and serum) were included for the untreated groups, and five biological replicates per tissue were included for the dietary intervention groups. Free fatty acid acids were performed only on brain samples.

Targeted lipidomic analyses were performed by the Columbia University Biomarkers Core Laboratory. Lipids were extracted using a modified Bligh and Dyer extraction protocol, and species were quantified using ultra-performance liquid chromatography-tandem mass spectrometry (UPLC-MS/MS). Lipid classes analyzed included phosphatidylcholines (PC), phosphatidylethanolamines (PE), sphingomyelins (SM), ceramides (Cer), lysophospholipids, and free fatty acids. Lipid values were normalized total protein content per sample. Statistical analysis was performed using one-way ANOVA to assess lipid abundance differences across experimental groups. Additionally, principal component analysis (PCA) was conducted separately for brain and serum samples to evaluate lipid shifts by genotype and drug treatment. All measurements and analytical procedures followed standardized protocols implemented by the Columbia Biomarkers Core.

### Glial engulfment assay

Postnatal day 5 (P5) Aldh1l1-eGFP::*Fmr1* wildtype and Aldh1l1-eGFP::*Fmr1* knockout, for astrocyte engulfment, and wildtype and *Fmr1* knockout, for microglial engulfment, as described.^19^ Mice were anesthetized by hypothermia and intraperitoneal injection of ketamine/xylazine, respectively. CTB conjugated to Alexa594 and Alexa 647 was intravitreally injected into the left and right eyes, as described in the RGC axon segregation assay, and allowed to recover for 48 hours. At P7, mice were transcardially perfused with 4% paraformaldehyde, and their heads fixed in 4% paraformaldehyde overnight. Brains were dissected, cryoprotected by serial incubation of 10% sucrose and 30% sucrose, and cryosectioned at 60 micron thickness throughout the dLGN.

Microglia were labeled immunohistochemically, using an antibody raised against the microglial marker P2RY12. The two most medial dLGN section were selected for analysis, and two field-of-view Z-stacks focusing on the ipsilateral and contralateral overlap zone were taken per section on a Nikon A1R confocal microscope, using the 63x oil objective (NA: 1.4, CFI60 Plan Apochromat). Images were processed in FIJI using the Remove Outlier (Radius 2.0 pixels, Threshold 20, Bright) and Subtract Background tools (Rolling ball radius: 50.0 pixels). Image analysis was performed using a custom Matlab program that quantifies the volume represented by the number of CTB-signal positive pixels that are both colocalized with and surrounded in all three dimensions Aldh1l1-eGFP-signal positive pixels.

## RESULTS

### Synaptic size and density show widespread decreases *Fmr1* knockout mice at P7

An overarching question in this study is whether synapse organization is disrupted in the retinogeniculate pathway in the *Fmr1* knockout model. To address this, synaptic puncta size and density were examined. Synapses were labeled using the presynaptic marker VGlut2 and the postsynaptic marker Homer in wild-type (WT) and *Fmr1* KO mice at postnatal day 7 (P7), a developmental stage where when retinogeniculate synapses are actively undergoing refinement (**Figure 1A-B**). Quantification revealed that synapse density was comparable between genotypes; however, average synapse size was significantly reduced in *Fmr1* KO dLGN compared to WT controls (**Figure 1C**). This difference was primarily driven by smaller postsynaptic puncta, consistent with prior reports of increased immature dendritic spines in other brain regions lacking FMRP.^45^ Additionally, presynaptic puncta in *Fmr1* KO mice trended smaller, but this difference was not statistically significant, suggesting that postsynaptic structures are more strongly affected at this developmental stage.

**Figure 1:**
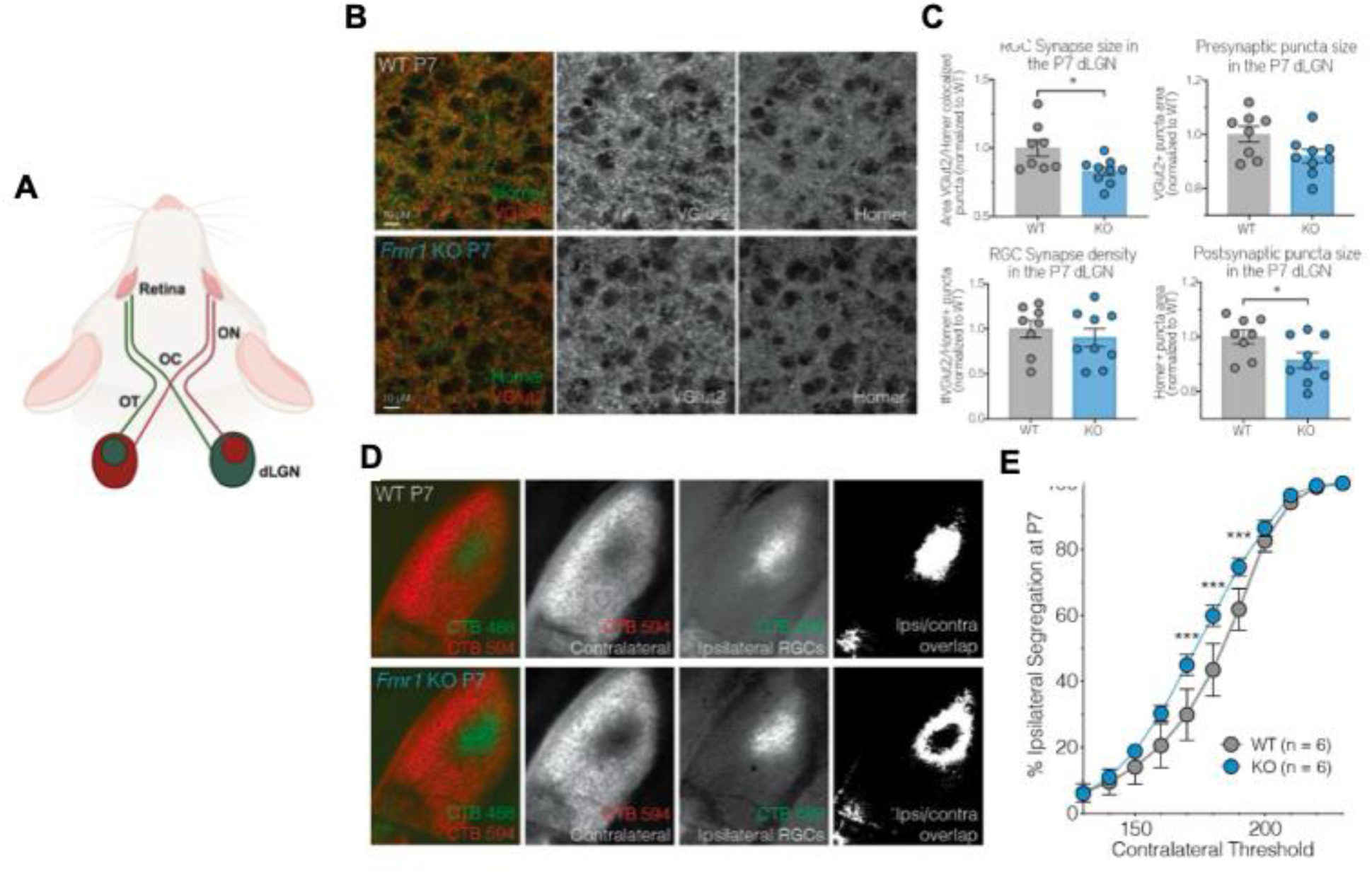
Altered synaptic architecture and increased retinal ganglion cell (RCG) segregation in the dorsal lateral geniculate nucleus (dLGN) of *Fmr1* KO mice at P7. **(A)** Schematic diagram showing the projection of RCG axons from the retina to the dLGN via the optic chiasm (OC), with ipsilateral (green) and contralateral (red) projections. **(B)** Field-of-view images from immunohistochemical labeling of presynaptic (VGlut2; red, center) and postsynaptic (Homer; green, right) puncta in the P7 dLGN of WT and *Fmr1* KO mice. *Images* were acquired using a Nikon A1R Confocal Microscope with a 63x oil objective. **(C)** Quantification of synaptic puncta using the ImageJ PunctaAnalyzer plug-in. The area of VGlut2+ and Homer+ colocalized puncta was significantly decreased in *Fmr1* KO mice (*p = 0.0285, Student’s t-test), whereas the size and number of VGlut2+ or Homer+ puncta alone were not significantly altered. **(D)** Representative images of ipsilateral and contralateral RGC inputs labeled via intravitreal injection of CTB-488 (green) and CTB-594 (red) in WT and *Fmr1* KO mice at P7. Grayscale images show segregated RGC inputs and overlap within the dLGN. **(E)** Quantification of ipsilateral segregation in the dLGN using a threshold-independent analysis. *Fmr1* KO mice exhibited significantly increased segregation compared to WT littermates (*p < 0.005, grouped analysis of multiple t-tests with Holm-Sidak correction for multiple comparisons, α = 5.00%).

To assess whether these structural synapse differences were accompanied by altered axonal refinement, eye-specific segregation of RGC inputs to the dLGN at P7 was examined. During normal development, RCG projections from both eyes initially overlap in the dLGN and are gradually pruned to form distinct ipsilateral and contralateral territories. Because refinement is actively ongoing at P7, this time point provides a sensitive window to detect differences in the timing or extent of segregation. Retinal inputs from each eye were labeled with spectrally distinct CTB fluorophores, and the degree of overlap in the dLGN was measured (**Figure 1D**). *Fmr1* KO mice exhibited significantly reduced overlap between ipsilateral and contralateral projections compared to WT mice, suggesting increased segregation at this developmental stage (**Figure 1E**). These results indicate that *Fmr1* KO mice exhibit both smaller synapses and increased segregation of RCG inputs at P7, pointing to early disruptions in the balance between synapse maturation and pruning in the thalamus.

### Cell-type-specific gene expression changes in postnatal day 7 (P7) *Fmr1* knockout mouse glia and neurons are associated with synaptic remodeling

Having identified altered retinogeniculate synapse development in the Fragile X syndrome (FXS) model, the next question raised was which cellular components mediate these changes during early postnatal development. To characterize the transcriptional changes associated with loss of *Fmr1,* single nucleus RNA sequencing (snRNAseq) was performed on dorsal lateral geniculate nucleus (dLGN) tissue of wildtype and *Fmr1* knockout mice at postnatal day 7 (P7), a critical developmental period marked by active synaptic refinement. Unsupervised clustering of the snRNAseq data using known cell-type markers revealed clear segregation of major neuronal and glial populations (**Figure S1**). Neurons formed the largest and most transcriptionally diverse population, while glial cells including astrocytes, microglia and oligodendrocytes formed well-defined clusters. Additionally, oligodendrocyte precursor cells (OPCs) showed separation from mature oligodendrocytes, reflecting distinct states within myelinating cell lineages. Endothelial cells and pericytes clustered together, representative of vascular tissue in this brain region. Gene expression data was then aggregated by individual mouse and cell-type to generate pseudobulk profiles for downstream analysis. This approach enabled robust statistical comparison across biological replicates by combining gene expression data at the individual mouse level, thereby reducing the technical noise inherent to snRNAseq data while maintaining cell-type-specific resolution. Pseudobulk differential expression analyses was performed for astrocytes, microglia, excitatory neurons and inhibitory neurons, as these cell-types are central to synaptic development, refinement, and maturation during early postnatal stages. Specifically, astrocytes and microglia are essential regulators of synapse remodeling as they engulf synapses in an activity-dependent manner.^46^ Neurons were further divided into excitatory and inhibitory populations to capture any differences in how *Fmr1* loss affects neuronal subtypes that exhibit distinct firing properties during development. Differential expression analysis was then performed for each cell-type. While this workflow identified many DEGs connected to synaptic pruning, it is important to note that many pathways associated with pruning are regulated via post-translational modifications, which may not be detected at the transcriptional level. Nonetheless, this approach enabled direct comparison of cell-type-specific transcriptional changes across glial and neuronal cells during a critical period of synaptic refinement.

Differential expression analysis of glial pseudobulk gene expression data identified several notable differentially expressed genes (DEGs) in *Fmr1* knockout mice compared to wildtype (WT) controls. Notably, many of the DEGs in microglia (**Figure 2A**, **Table 1**) and astrocytes (**Figure 2C**) were linked to synaptic pruning and remodeling, processes that have implicated in FXS.^47^ In microglia, loss of *Fmr1* resulted in the downregulation of *Map1b*, Tub2b, and *Tuba1a* (**Figure 2B**, **Table 2**), two microtubule associated genes that are important for neuronal circuit development.^48,49^ Additionally, the enzymatic severing of microtubules is known to occur early in the pruning process,^50^ and the post-translational modification tubulin tags microtubules for severing,^51^ making these genes important regulators of processes needed for pruning. The actin genes, *Actb* and *Actg1* were also differentially expressed in astrocytes; this is intriguing, as actin is essential in phagocytotic cup formation, which is required for engulfment.^52^ Finally, *Sparc* was upregulated, which has been shown to promote cell-autonomous synapse elimination.^53^ Consistent with these findings, gene ontology (GO) enrichment analysis of downregulated microglia genes revealed significant enrichment for pathways related to synaptic vesicle endocytosis and microtubule dynamics (**Supplementary Figure S2**).

**Figure 2:**
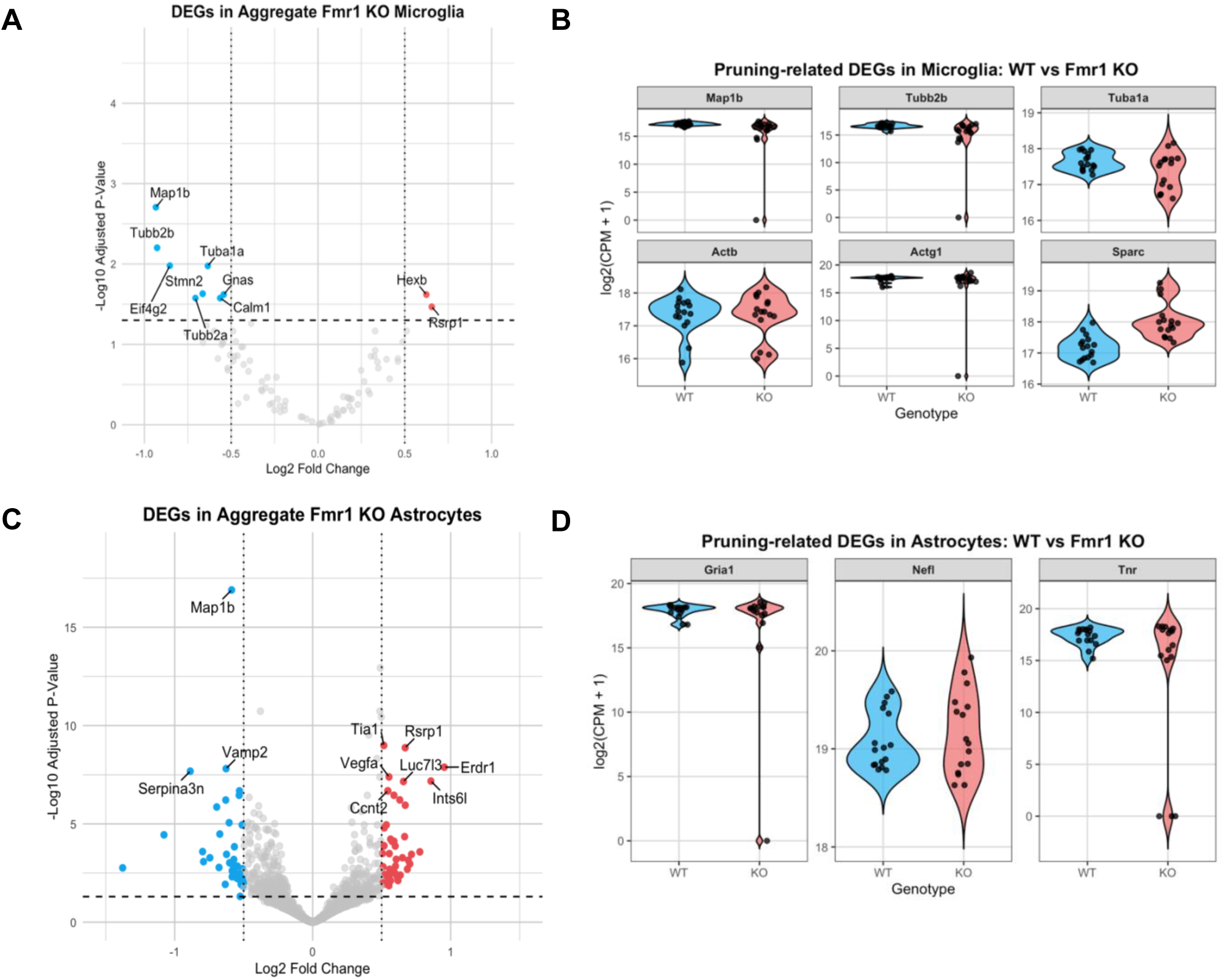
*Fmr1* deletion alters glial transcriptional programs, disrupting genes linked to synaptic pruning. **(A)** Volcano plot of differentially expressed genes (DEGs) in pseudobulked microglia from *Fmr1* KO versus WT mice. Genes significantly upregulated (red) or downregulated (blue) were defined using an adjusted p-value < 0.05 and |log₂FC| ≥ 0.25. **(B)** Violin plots showing differentially expressed genes of genes linked to the pruning process in microglia including the upregulated *Map1b, Tub2b, Tub1a*, *Act1b*, *Actg1*, and the upregulated gene *Sparc*. **(C)** Volcano plot of DEGs in pseudobulked astrocytes from *Fmr1* KO versus WT mice, with significantly up- and downregulated genes highlighted. (D) Violin plots showing reduced expression of synaptic and microtubule-associated genes including the downregulated genes *Gria1, Nefl,* and *Tnr*.

**Table 1:**
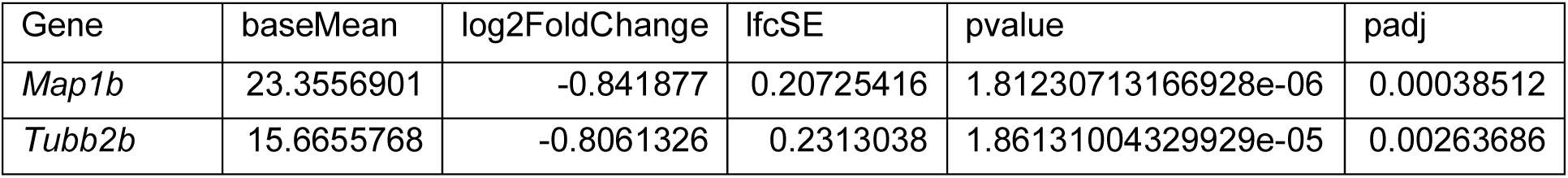
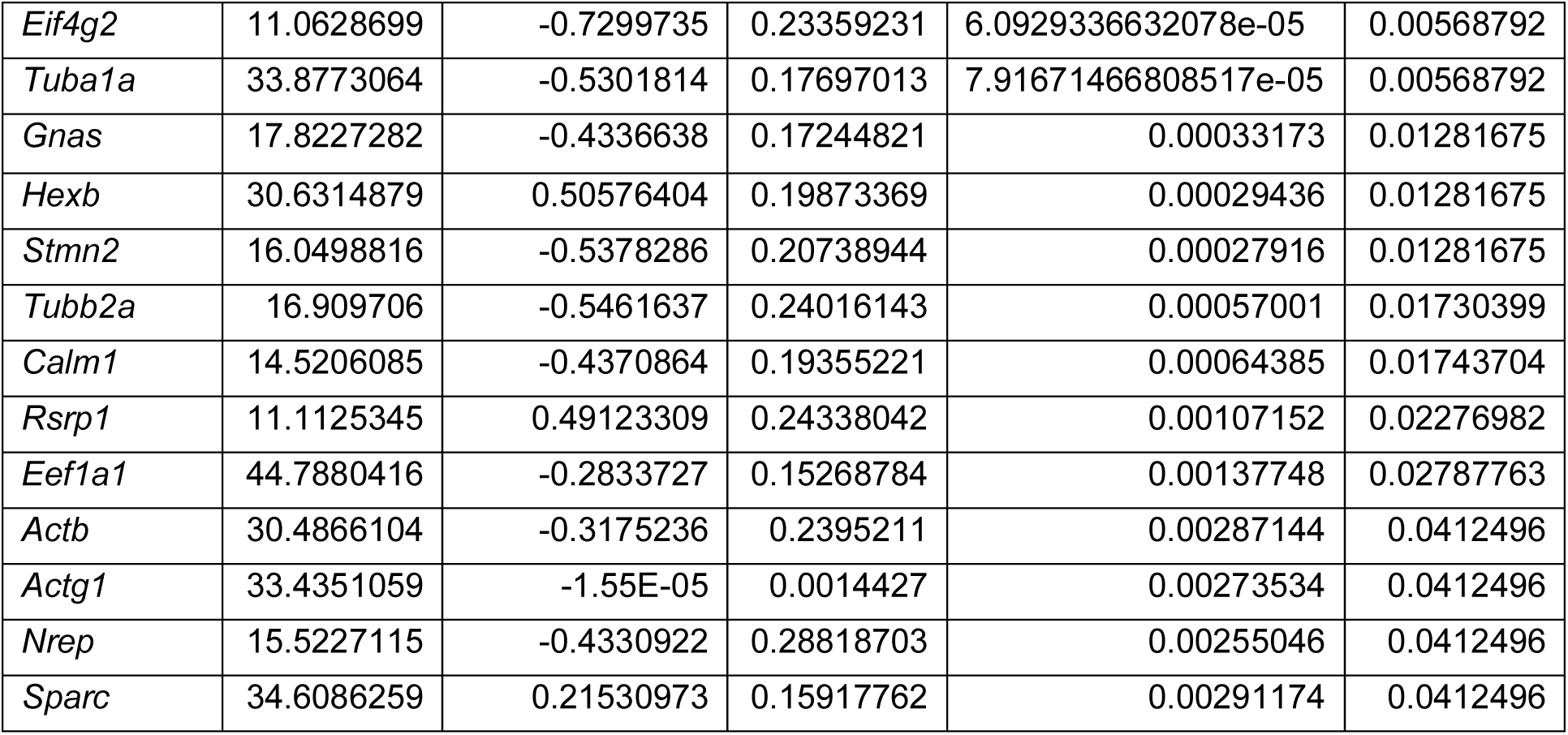
Top 15 differentially expressed genes in aggregated microglia from the dorsal lateral geniculate nucleus of *Fmr1* knockout (KO) mice compared to wildtype controls. Genes are ranked by statistical significance (adjusted p-value) and have a base mean above 10. Positive and negative log2FoldChange values indicate upregulation and downregulation, respectively, in *Fmr1* KO microglia.

**Table 2:**
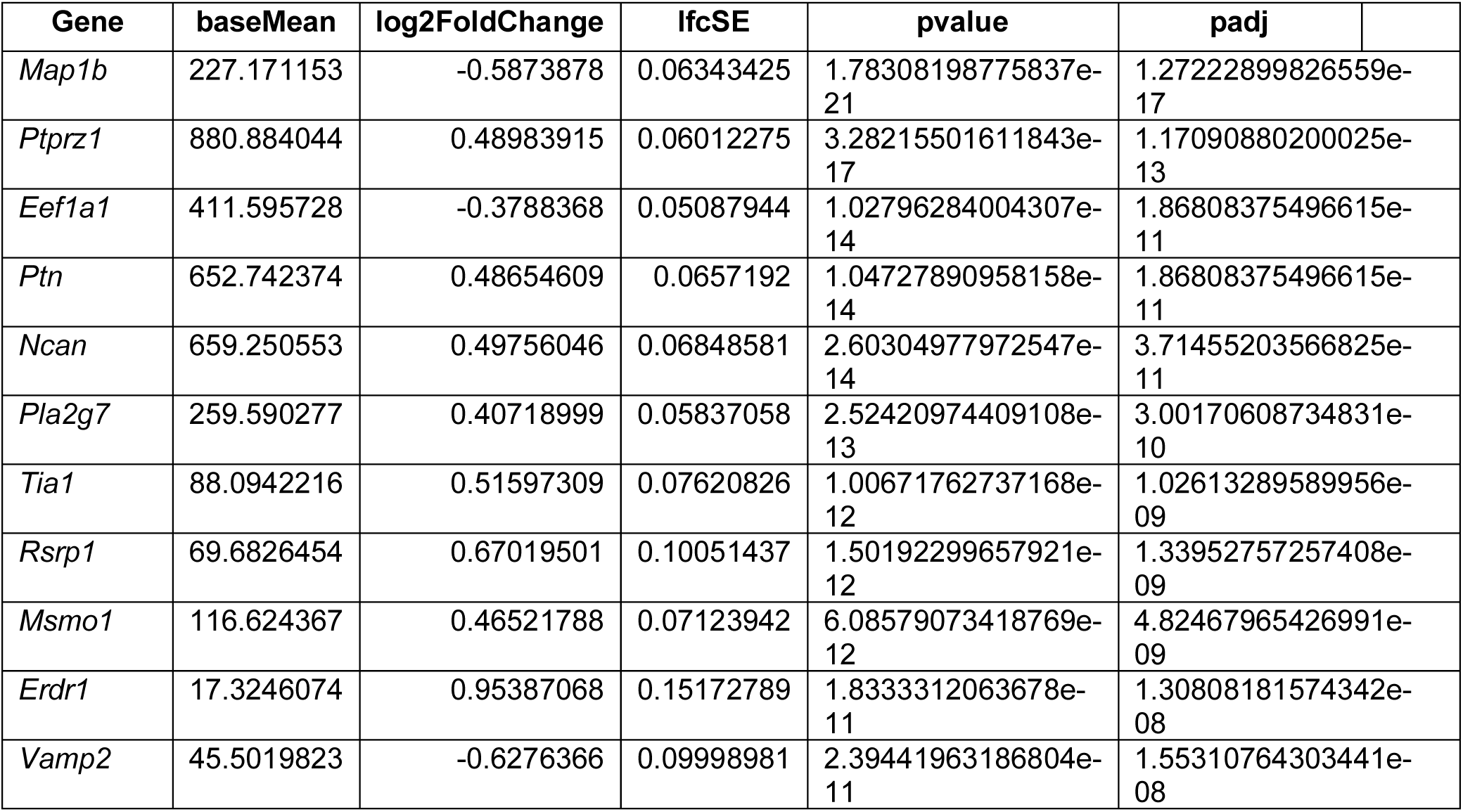
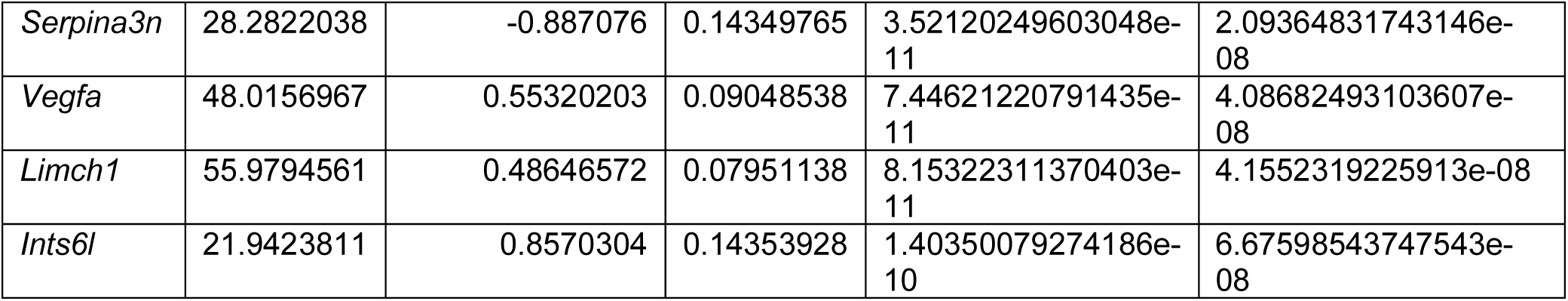
Top 15 differentially expressed genes in aggregated astrocytes from the dorsal lateral geniculate nucleus of *Fmr1* knockout (KO) mice compared to wildtype controls. Genes are ranked by statistical significance (adjusted p-value) and have a base mean above 10. Positive and negative log2FoldChange values indicate upregulation and downregulation, respectively, in *Fmr1* KO astrocytes.

In astrocytes, loss of *Fmr1* was associated with the downregulation of synaptic stabilization genes including *Gria1, Nefl, and Tnr* (**Figure 2D**). This may reflect a transcriptional shift in astrocytes from a synapse-supportive state, to one that favors remodeling or pruning. As in microglia, *Map1b* was downregulated in astrocytes, indicative of altered microtubule remodeling pathways across both glial cell-types. GO enrichment analysis of astrocytic DEGs further demonstrated that downregulated genes were enriched for synaptic and translational processes, while upregulated genes were enriched for fibroblast growth factor receptor apoptotic signaling, pointing toward altered regulation of pathways related to synaptic cell survival (**Figure S3**). Together, these results suggest that the absence of *Fmr1* is accompanied by coordinated, cell-type-specific alterations in glial gene expression, potentially affecting synaptic remodeling dynamics during development.

Next, pseudobulk differential expression analysis in neurons was completed. In excitatory neurons, many downregulated DEGs were associated with synaptic organization, extracellular matrix composition, and activity-dependent remodeling. Notably, genes such as *Bcan, Ptprz1, Sparc, Sparc1, Mfge8*, and *Tnr* showed reduced expression in *Fmr1* knockout mice (**Figure 3A-B**, **Table 3**), several of which encode proteins that help regulate synaptic stability and mark synapses for elimination. For example, *Bcan* and *Tnr* are core components of the perisynaptic extracellular matrix and function to maintain synaptic integrity,^54^ while *Mfge8* is a neuroprotective glycoprotein that facilitates the clearance of apoptotic cells, and *Ptprz1* is a known promoter of synaptic formation.^55^ Notably, *Map1b* and *Sparc* were also downregulated in excitatory neurons as in glia, further supporting disrupted cytoskeletal remodeling and synapse elimination. Together, these DEGs are consistent with altered gene expression signatures associated with synaptic structure and maturation, mechanisms that determine pruning potential. GO analysis of downregulated genes in excitatory neurons further revealed enrichment of pathways related to cytoplasmic and presynaptic translation, neuron fate specification, and neurofilament bundle assembly (**Figure S4**). Thus, loss of *Fmr1* may shift excitatory neurons toward a transcriptional state associated with reduced synaptic stability and enhanced pruning susceptibility.

**Figure 3:**
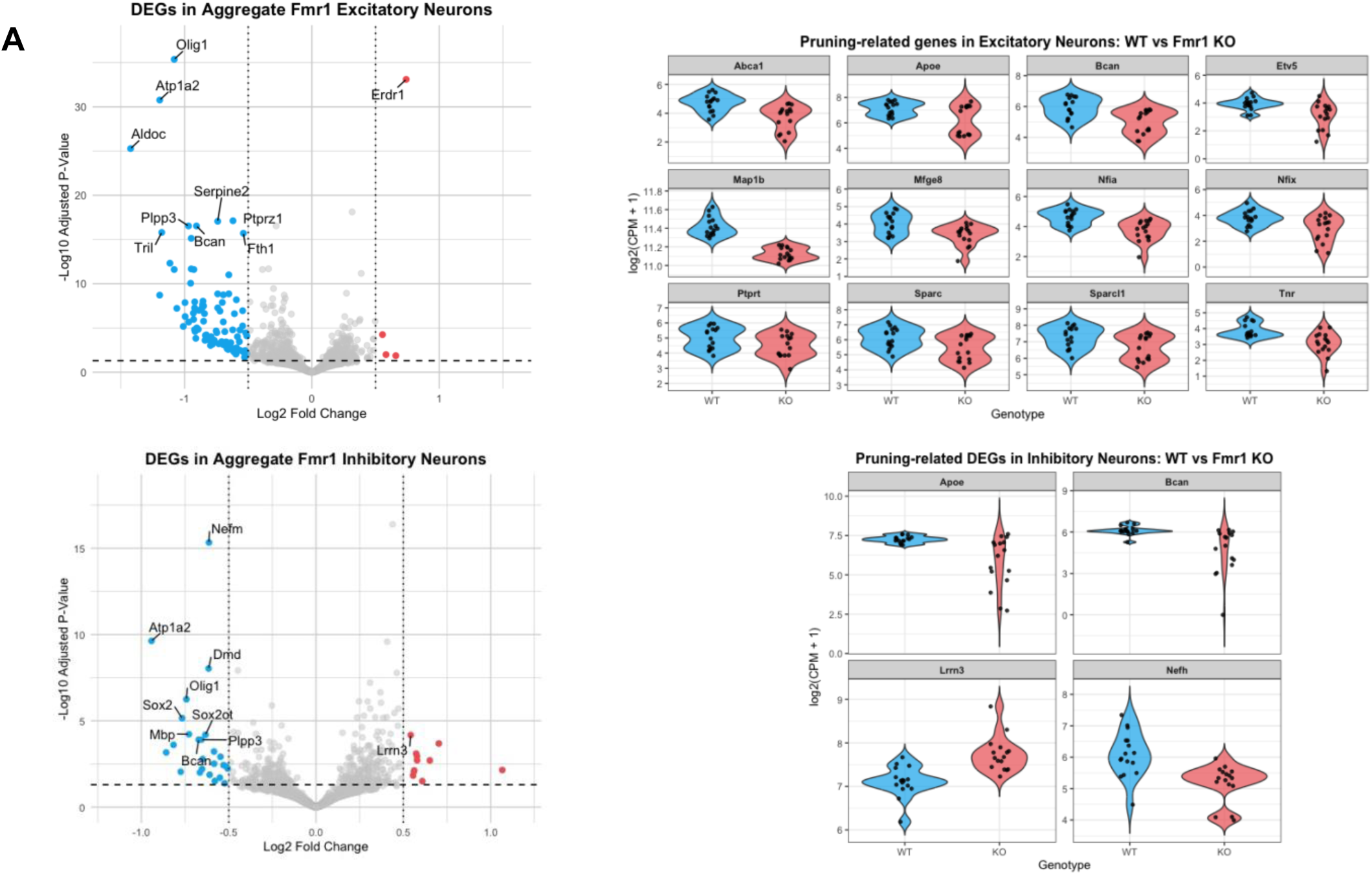
*Fmr1* deletion disrupts genes linked to synaptic pruning in excitatory and inhibitory neurons in the P7 *Fmr1* KO dLGN. **(A)** Volcano plot of differentially expressed genes (DEGs) in pseudobulked excitatory neurons from *Fmr1* KO versus WT mice. Genes significantly upregulated (red) or downregulated (blue) were defined as having adjusted p-value < 0.05 and |log₂FC| ≥ 0.25. Several downregulated genes are associated with synaptic organization, extracellular matrix structure, and activity-dependent remodeling. **(B)** Violin plots showing reduced expression of pruning-and matrix-associated genes *including Bcan, Ptprt, Sparc, Sparc1, Map1b, Mfge8, Nfia, Nfix, Apoe, Etv5, and Tnr* in excitatory neurons from *Fmr1* KO mice compared to WT littermates. **(C)** Volcano plot of DEGs in pseudobulked inhibitory neurons from *Fmr1* KO versus WT mice, highlighting significantly up- and downregulated transcripts. **(D)** Violin plots displaying altered expression of pruning- and cytoskeleton-related genes in inhibitory neurons, including *Apoe, Bcan, Lrrn3,* and *Nefh*, demonstrating that transcriptional signatures associated with synaptic stability and remodeling are disrupted across both major neuronal populations in the P7 *Fmr1* KO dLGN.

**Table 3:**
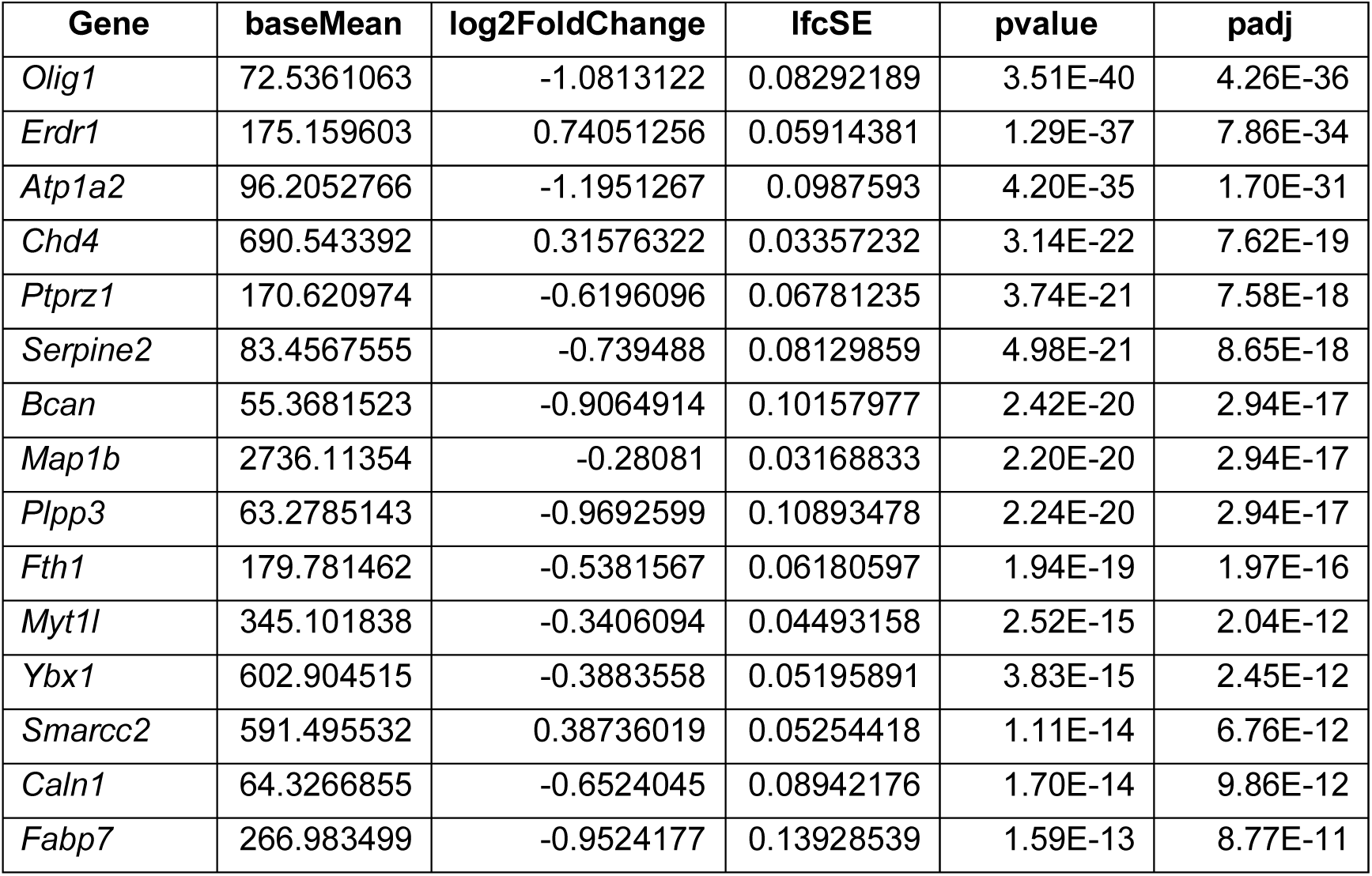
Top 15 differentially expressed genes in aggregated excitatory neurons from the dorsal lateral geniculate nucleus of *Fmr1* knockout (KO) mice compared to wildtype controls. Genes are ranked by statistical significance (adjusted p-value) and have a base mean above 50. Positive and negative log2FoldChange values indicate upregulation and downregulation, respectively, in *Fmr1* KO excitatory neurons.

| Gene | baseMean | log2FoldChange | lfcSE | pvalue | padj |
| --- | --- | --- | --- | --- | --- |
| *Olig1* | 72.5361063 | -1.0813122 | 0.08292189 | 3.51E-40 | 4.26E-36 |
| *Erdr1* | 175.159603 | 0.74051256 | 0.05914381 | 1.29E-37 | 7.86E-34 |
| *Atp1a2* | 96.2052766 | -1.1951267 | 0.0987593 | 4.20E-35 | 1.70E-31 |
| *Chd4* | 690.543392 | 0.31576322 | 0.03357232 | 3.14E-22 | 7.62E-19 |
| *Ptprz1* | 170.620974 | -0.6196096 | 0.06781235 | 3.74E-21 | 7.58E-18 |
| *Serpine2* | 83.4567555 | -0.739488 | 0.08129859 | 4.98E-21 | 8.65E-18 |
| *Bcan* | 55.3681523 | -0.9064914 | 0.10157977 | 2.42E-20 | 2.94E-17 |
| *Map1b* | 2736.11354 | -0.28081 | 0.03168833 | 2.20E-20 | 2.94E-17 |
| *Plpp3* | 63.2785143 | -0.9692599 | 0.10893478 | 2.24E-20 | 2.94E-17 |
| *Fth1* | 179.781462 | -0.5381567 | 0.06180597 | 1.94E-19 | 1.97E-16 |
| *Myt1l* | 345.101838 | -0.3406094 | 0.04493158 | 2.52E-15 | 2.04E-12 |
| *Ybx1* | 602.904515 | -0.3883558 | 0.05195891 | 3.83E-15 | 2.45E-12 |
| *Smarcc2* | 591.495532 | 0.38736019 | 0.05254418 | 1.11E-14 | 6.76E-12 |
| *Caln1* | 64.3266855 | -0.6524045 | 0.08942176 | 1.70E-14 | 9.86E-12 |
| *Fabp7* | 266.983499 | -0.9524177 | 0.13928539 | 1.59E-13 | 8.77E-11 |

Inhibitory neurons also exhibited genotype-dependent transcriptional changes. *Bcan* was downregulated in inhibitory neurons in addition to excitatory neurons, indicating a shared reduction in synapse structure across neurons, and supporting a shift toward a more plastic, pruning-permissive synaptic environment. Other downregulated genes include *ApoE*, *Lrrn3*, and *Nefh* (**Figure 3C-D**, **Table 4**). *ApoE* has been shown to control levels of the complement protein C1q, thereby regulating the rate of synaptic pruning by astrocytes.^18^ Further, *Lrrn3* encodes a transmembrane protein that inhibits glycolysis to reduce mitochondrial dysfunction and cell apoptosis,^56^ suggesting that its reduced expression may increase neuronal vulnerability to pruning.^57^ In agreement with this, GO enrichment analysis of inhibitory neuron DEGs revealed a downregulation of pathways associated with cytoskeletal organization and axon-related processes (**Figure S5**). These results highlight changes in cellular structural and trafficking mechanisms in inhibitory neurons that may influence synaptic stability and susceptibility to pruning and remodeling.

**Table 4:**
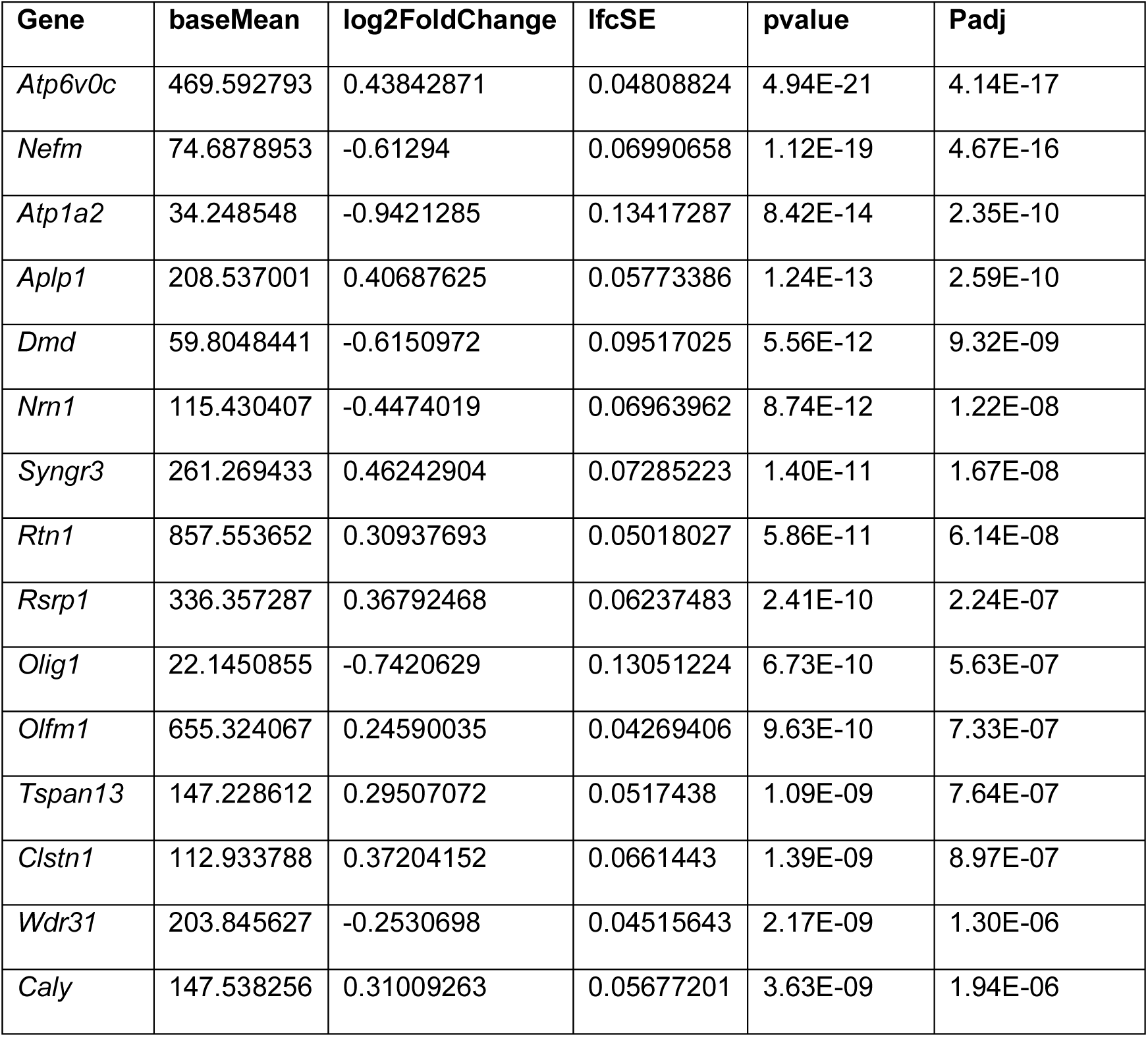
Top 15 differentially expressed genes in aggregated inhibitory neurons from the dorsal lateral geniculate nucleus of *Fmr1* knockout (KO) mice compared to wildtype controls. Genes are ranked by statistical significance (adjusted p-value) and have a base mean over 10. Positive and negative log2FoldChange values indicate upregulation and downregulation, respectively, in *Fmr1* KO inhibitory neurons.

### Astrocyte-to-microglia signaling is likely increased in *Fmr1* KO mice via multiple lipid pathways

The single-cell transcriptomics data identified relevant pathway changes in astrocytes, microglia, and both classes of neurons, but it is unclear from DEG analysis alone whether there are mutual interactions amongst the different cell-types. Given that signaling between neurons and glia is an essential aspect of activity-dependent pruning that is tightly regulated, the potential interactions between these cell-types were further investigated.

Cell-Chat, a computational tool that infers intercellular signaling from known ligand-receptor interactions using gene expression data,^41^ was applied to pseudobulk snRNAseq profiles from the mouse dLGN. This analysis enabled direct comparison of cell-cell signaling networks between WT and *Fmr1* KO mice, revealing specific ligand-receptor pathways with altered activity in the absence of FMRP. This modeling revealed that glial communication networks were markedly altered in *Fmr1* KO mice, with an increase in overall signaling activity relative to WT controls. The most pronounced change was an enhancement of astrocyte-to-microglia signaling (**Figure 4A**), suggesting increased astrocytic output and heightened microglial responsiveness.

**Figure 4:**
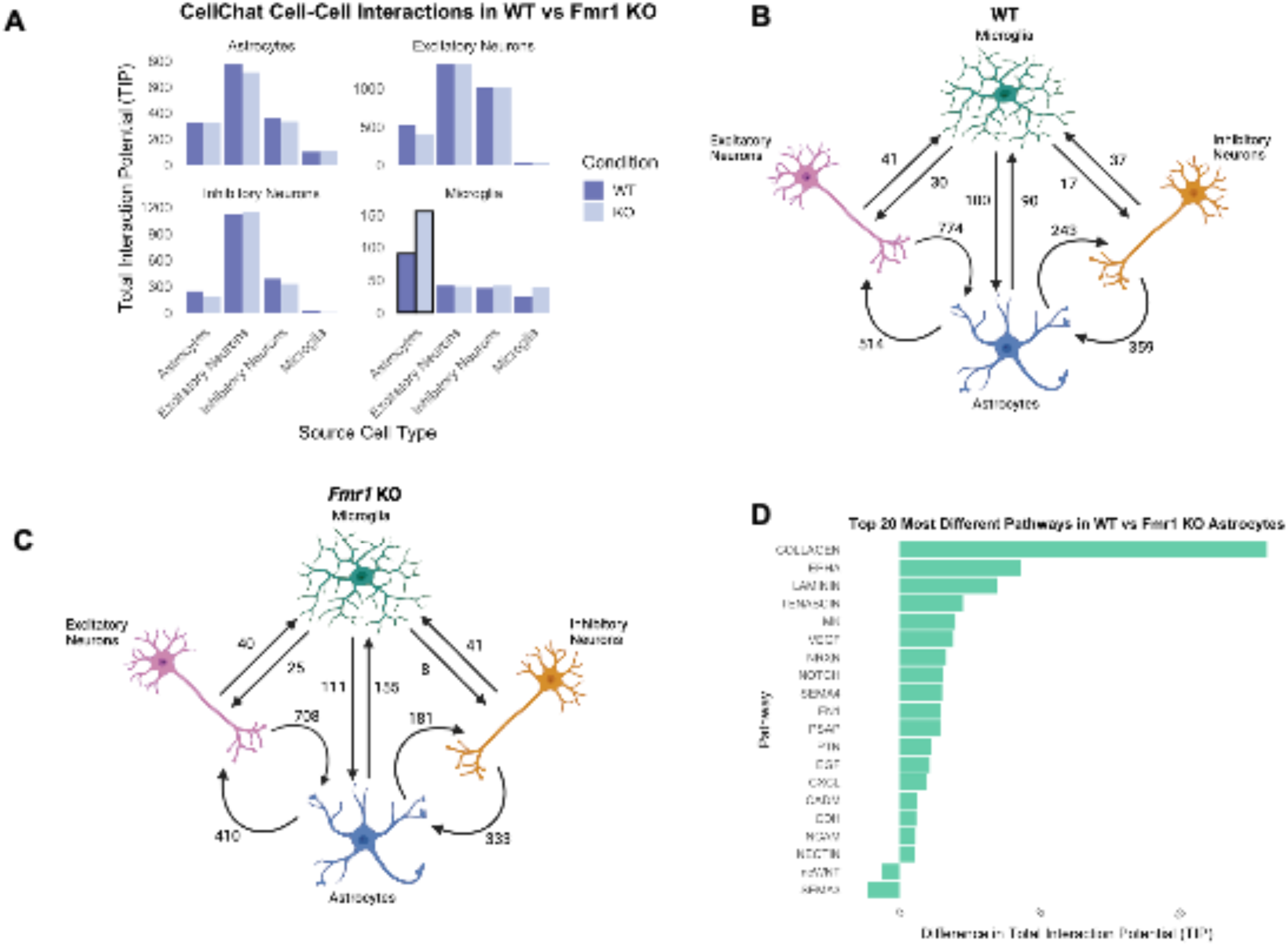
Astrocyte-to-microglia signaling is increased in *Fmr1* knockout (KO) mice and enriched for synaptic lipid-associated pathways. **(A)** Total interaction potential (TIP) scores derived from CellChat analysis showing overall cell-cell communication in wild-type (WT) and *Fmr1* KO dorsal lateral geniculate nucleus (dLGN), stratified by source and target cell-types. Astrocyte-to-microglia signaling is increased in the KO condition. **(B-C)** CellChat signaling network diagrams illustrating the number and strength, TIP score, of predicted ligand-receptor interactions between glial and neuronal populations in WT **(B)** and *Fmr1* KO **(C)** mice. Astrocytes in the KO exhibit elevated outgoing signals to microglia. **(D)** Top 20 pathways with the greatest difference in TIP between WT and KO astrocytes. Several lipid- and synapse-related pathways are enriched in the KO, including ephrin type a (EPHA) and semaphorin (SEMA) signaling.

To quantify this observation, a custom metric called Total Interaction Potential (TIP) was used, calculated as the sum of ligand-receptor interactions inferred between sender and receiver cell-types across all signaling pathways. When applied to signals originating from astrocytes, TIP was substantially increased for astrocyte-to-microglia signaling in *Fmr1* KO mice (**Figure 4B-C**). This quantification confirmed that astrocytes are the predominant source of altered glial signaling upon loss of FMRP and provided a framework for identifying the ligand-receptor pairs and signaling pathways that drive that changes. Individual pathway level contributions to TIP were next examined to identify which signaling mechanisms drive the increased astrocyte-to-microglia communication in *Fmr1* KO mice. The absolute TIP was increased for Ephrin type-A receptor signaling (EphA) and semaphorin (SEMA) signaling, which are both known regulators of synaptic connectivity and glial function. EphA signaling, mediated by ephrin receptor tyrosine kinases and their ephrin ligands, has a critical role in dendritic spine morphogenesis, synapse formation, and synaptic plasticity. Specifically, EphA signaling facilitates synaptic maturation by regulating spine structure and cytoskeletal dynamics.^58^ These interactions also influence synaptic architecture by organizing lipid rafts, specialized membrane domains enriched with signaling molecules that affect synapse formation and stability. Interestingly, loss of ephrin-A2 has previously been shown to enhance experience-dependent synaptic pruning.^59^ Thus, the increased EphA pathway activity observed in *Fmr1* knockout mice may reflect an attempted compensatory mechanism by glial cells to suppress synaptic pruning during a period of heightened remodeling.

In parallel, the SEMA pathway, particularly SEMA3A (semaphorin 3A) is critical in axon guidance, dendritic patterning, and synaptic refinement during development. Through interactions with neuropilin and plexin receptors, SEMA3A influences cytoskeletal remodeling by activating downstream pathways that drive growth cone collapse and synaptic pruning, processes essential for neural circuit refinement. In addition to its role in neural wiring, SEMA signaling also affects glial function and neuron-glia interactions. A recent study showed that SEMA3A triggers the degradation of FMRP in hippocampal neural growth cones via the ubiquitin-proteasome pathway, prompting translational repression of targets such as MAP1B.^60^ Consistent with this mechanism, *Map1B* was identified as downregulated in both astrocytes and microglia (**Figure 2**), reinforcing the pathological relevance of disrupted SEMA3A-FMRP-Map1B signaling. This coordinated downregulation across cell-types suggests that this pathway may drive cytoskeletal instability in relation to synaptic pruning, and ultimately synaptic dysfunction, in FXS.

### Preliminary lipidomic profiling implicates altered EphA-associated lipids in *Fmr1* KO mice, which may be reversed with Lovastatin

Given that EphA and SEMA lipid-associated signaling pathways emerged as top contributors to aberrant astrocyte-microglia interactions in *Fmr1* KO mice, exploratory lipidomic profiling was performed to determine whether loss of FMRP alters the abundance of lipid species essential for neuronal development. This approach was motivated by the established requirement of both EphA and SEMA receptors for precise membrane microenvironments, particularly in lipid rafts which are enriched in cholesterol, sphingolipids and phospholipids.^61,62^ Moreover, the disruption of lipid domains can impair receptor localization, clustering, and signaling involved in synaptic development and glial communication.^63^

Lipidomic analysis of brain and serum from *Fmr1* KO mice revealed significant reductions in several EphA-associated lipid species, including phosphatidylcholines (PC 38:4, PC 38:6), phosphatidylethanolamine (PE 36:2), and sphingomyelins (SM d18:1/22:0, SM d18:1/20:0), compared to WT controls (**Figure S6A, C**). These lipid classes are known components of membrane microdomains such as lipid rafts that can support EphA receptor localization and signaling at the synapse. However, the observed changes were small in magnitude and exploratory in nature. Therefore, these data only suggest a potential association between FMRP loss and lipid dysregulation. The increased EphA pathway activity predicted by CellChat may reflect a compensatory transcriptional response to membrane stability; however, without direct assessment of receptor localization or signaling, this interpretation remains speculative in nature.

To explore whether the lipid changes observed in *Fmr1* KO mice are modifiable, mice were treated with the cholesterol lowering drug Lovastatin via daily intraperitoneal injections from P4 to P6, and lipid levels were reassessed at P7. Cholesterol has been shown to decrease assembly and activation of EphA receptors,^64^ so it was presumed that Lovastatin would relieve this inhibition and enhance EphA levels. Indeed, this intervention led to a restoration of several EphA-associated lipid species, most notably PC 38:4 and PC 38:6, in both brain and serum of *Fmr1* KO mice (**Figure S6B, D**). However, the absence of a vehicle control and the modest magnitude of these effects limit interpretation, and these data do not establish a direct effect on Lovastatin on EphA receptor localization, signaling, or synaptic function. As such, these findings should be considered preliminary and hypothesis-generating, suggesting that membrane lipid composition may be pharmacologically sensitive in the context of FMRP loss. Additional validation will be required to determine whether lipid modulation meaningfully contributes to the phenotypes associated with FXS.

### Microglia and astrocytes over-engulf retinal ganglion cell synapses in *Fmr1* KO mice

The increased eye-specific segregation and smaller synapse size observed in the *Fmr1* KO dLGN indicate that synaptic pruning may occur either earlier or more extensively than in normal early development. In line with this, differential expression analysis of pseudobulk snRNAseq from *Fmr1* KO showed that DEGS in astrocytes and microglia were related to synaptic modeling and pruning, and cell-cell interaction analysis also implicated these two cell types as being potentially dysregulated together (**Figures 1-4**). Since microglia are known sculptors of synapses in the developing dLGN and inhibition of microglia phagocytosis reduces segregation of ipsilateral and contralateral retinal inputs,^19^ microglial engulfment of RGC synapses was examined to determine whether it is enhanced in the *Fmr1* KO dLGN.

To directly explore this suggested cellular activity, glial engulfment of retinal inputs was examined in *Fmr1* KO mice. RGC axons were labeled from each eye and the volume of presynaptic material contained within microglia or astrocytes in the P7 dLGN was quantified. To assay phagocytic activity, RCGs were labeled via intraocular injection of cholera toxin B (CTB) conjugated to Alexa Fluor-594 and Alexa Fluor-647 at P5, and mice were sacrificed at P7. Microglia were labeled via immunostaining with P2RY12, and astrocytes were identified using the Aldh1l1-eGFP genetic label. Confocal imaging was used to quantify engulfment as the volume of CTB-labeled presynaptic material contained within the P2RY12 microglia and Aldh1l1-eGFP astrocytes, normalized to total cell volume.

Microglia engulfment of RGC inputs was significantly increased in *Fmr1* KO mice compared to wild-type littermates (**Figure 5A-C**), although total microglial volume remained unchanged. This indicates that microglia are more actively phagocytosing retinal inputs in the absence of FMRP, rather than undergoing general changes in cell size or proliferation. Astrocytes also displayed a significant increase in engulfed RGC input volume in *Fmr1* KO mice (**Figure 5D-F**). As with microglia, this was not accompanied by a change in total astrocyte volume. This suggests that the over-engulfment phenotype is not restricted to microglia but involves both major glial populations known to contribute to synaptic pruning in the dLGN. Together with the multi-omics analyses, these findings demonstrate that both microglia and astrocytes exhibit elevated engulfment of retinal inputs in the developing *Fmr1* KO dLGN. Moreover, this elevated phagocytic activity across both microglia and astrocytes likely drives the reduction in synapse size and the accelerated eye-specific segregation observed in P7 *Fmr1* KO mice.

**Figure 5:**
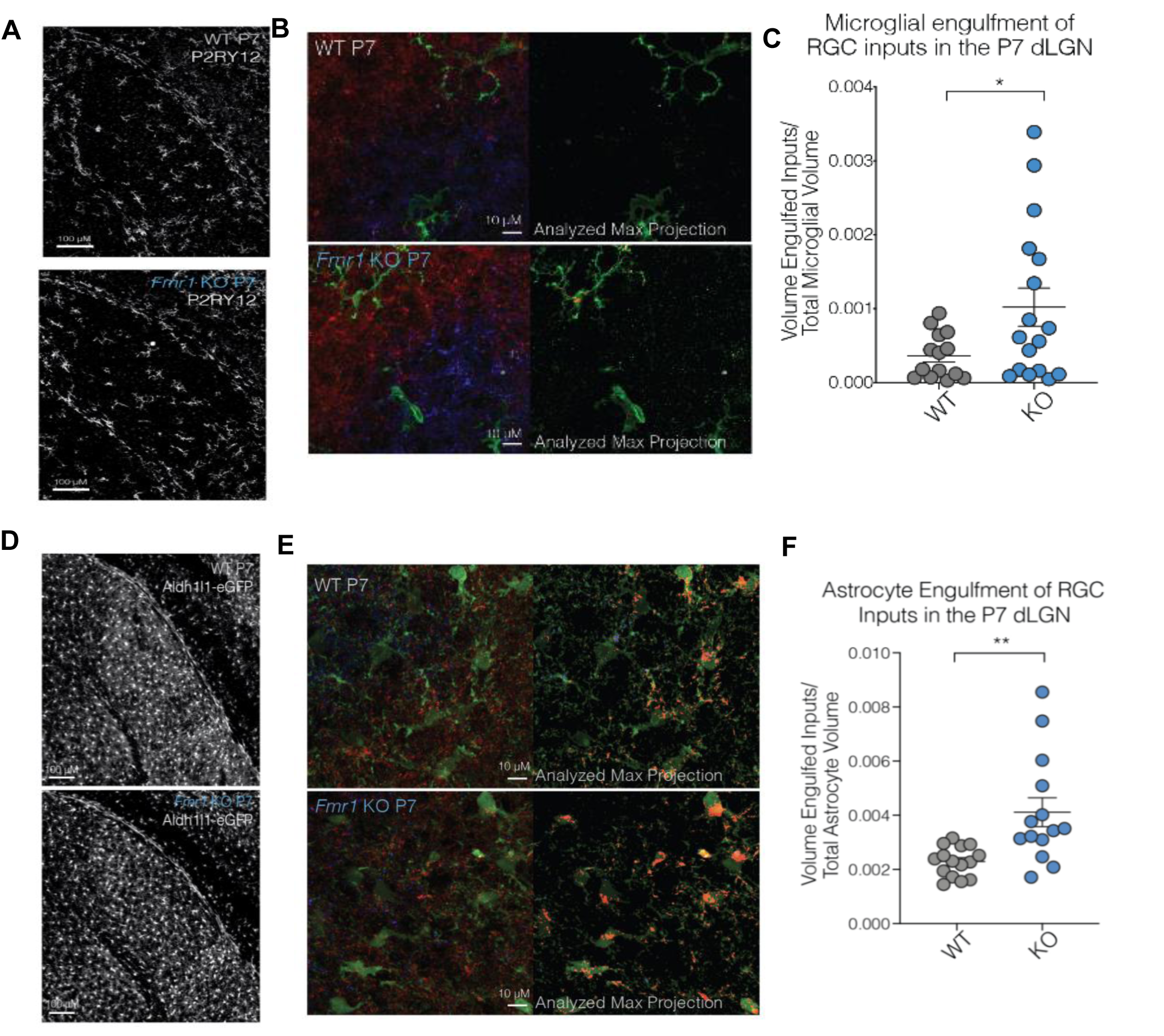
Microglia and astrocytes over-engulf retinal ganglion cell synapses in *Fmr1* KO mice. **(A)** P2RY12 immunolabeling reveals microglia in the dLGN of P7 WT and *Fmr1* KO mice. **(B)** Representative field-of-view z-stacks for microglial engulfment assay, with microglia labeled by P2RY12 (green), ipsilateral RGCs labeled by CTB-647 (blue), and contralateral RGCs labeled by CTB-594 (red). Maximum intensity projections of raw confocal images (left) and volume-quantified outputs (right) show engulfed RGC input within microglia. **(C)** Quantification of engulfed RGC inputs normalized to total microglial volume reveals significantly increased microglial engulfment in *Fmr1* KO mice compared to WT littermates (*p < 0.05, unpaired t-test; each dot represents one animal). **(D)** Aldh1l1-eGFP mice show transgenic labeling of astrocytes in the dLGN of P7 WT and *Fmr1* KO mice. **(E)** Representative field-of-view z-stacks for astrocyte engulfment assay, with astrocytes labeled by eGFP (green), ipsilateral RGCs labeled by CTB-647 (blue), and contralateral RGCs labeled by CTB-594 (red). Maximum intensity projections of raw confocal images (left) and volume-quantified outputs (right) show engulfed RGC input within astrocytes. **(F)** Quantification of engulfed RGC inputs normalized to total astrocyte volume reveals significantly increased astrocytic engulfment in *Fmr1* KO mice compared to WT littermates (**p = 0.0049, unpaired t-test with Welch’s correction; each dot represents one animal).

## Discussion

Fragile X syndrome is characterized by widespread molecular, cellular and circuit-level abnormalities that emerge during early brain development. In this study, an integrative multi-omic approach was applied to *Fmr1* KO mice at P7, a critical developmental stage for synaptic development. P7 *Fmr1* KO mice exhibited smaller synapse size and increased eye-specific segregation. This was accompanied by transcriptomic profiling that revealed dysregulation of genes involved in synaptic remodeling and glial engulfment. CellChat analysis showed enhanced astrocyte-to-microglia communication through EphA and SEMA pathways in FXS mice. Preliminary lipidomic analysis implicated disruption in EphA-associated membrane lipids that support receptor signaling within lipid rafts that may be reversed by Lovastatin. Finally, *Fmr1* KO mice showed increased glial synaptic engulfment upon loss of *Fmr1* in the dLGN. Together, these complementary approaches reveal coordinated molecular, cellular, and synaptic alterations that may underlie the pathogenesis of FXS.

To summarize these findings, a hierarchical model was constructed (**Figure 6**) illustrating how molecular disruptions cascade into higher-order cellular and structural abnormalities in P7 *Fmr1* KO mice. At the molecular level, exploratory lipidomic profiling revealed reductions in EphA-associated lipids PC 38:4 and PC 38:6 in both brain and serum, which are critical components of lipid rafts required for receptor localization and signaling. However, due to the limited strength of this experiment, this step in the model remains tentative and is denoted with a question mark, reflecting the uncertainty regarding the functional impact of these lipid changes. CellChat cell-cell communication analysis predicted increased astrocyte-to-microglia communication, as reported for synapse pruning during normal development,^46,65^ and here in FXS via the EphA pathway. These changes were accompanied by increased microglial and astrocytic engulfment of retinal ganglion cell (RGC) synapses, reduced postsynaptic puncta size, and increased eye-specific segregation. Ultimately, these phenotypes are triggered by transcriptional changes induced by loss of FMRP. Together, this model highlights a coordinated cascade in which differential gene expression leads to lipid imbalances, which alter glial communication to drive cellular and synaptic changes that ultimately reshape developing circuits in the absence of FMRP.

**Figure 6:**
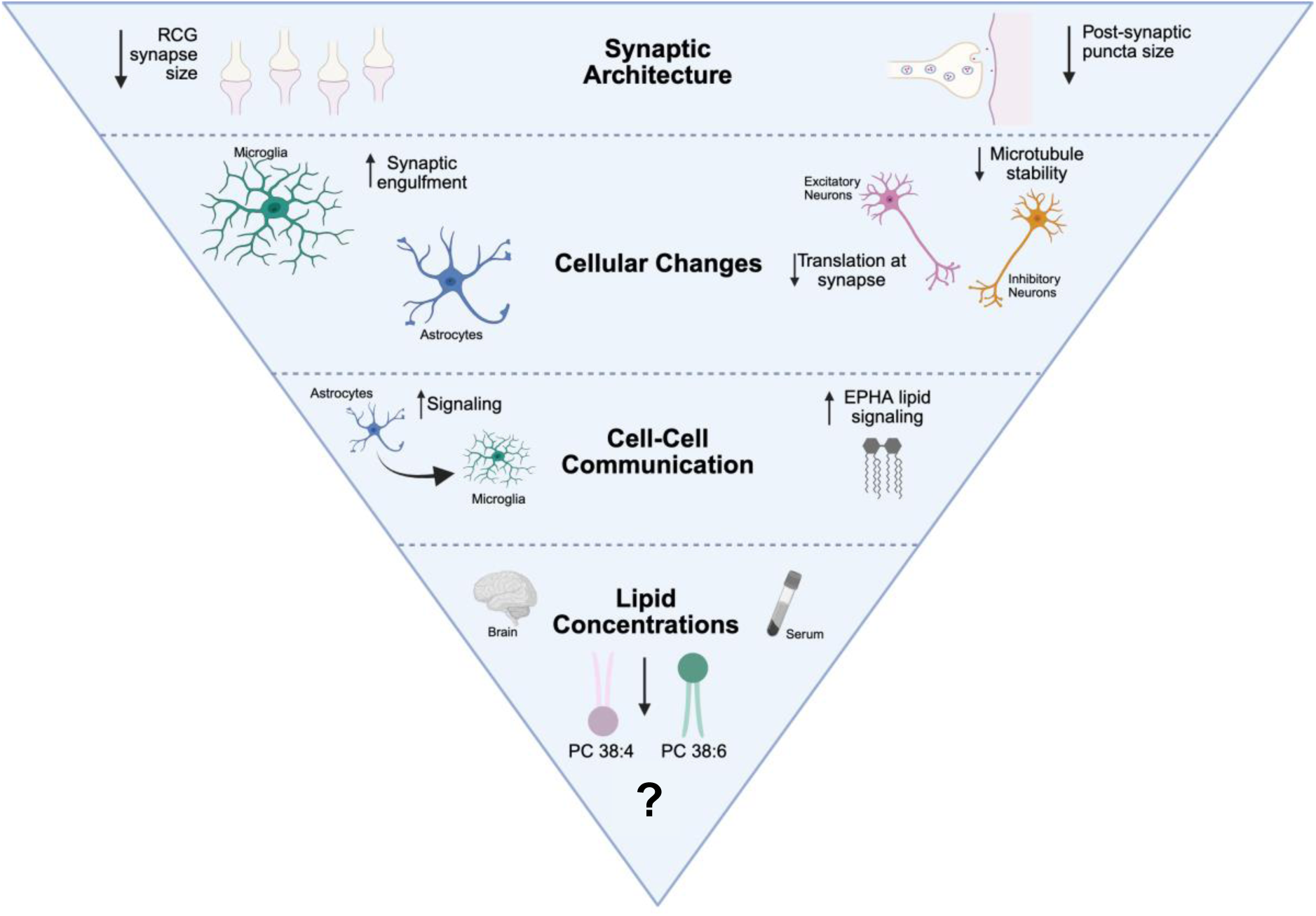
Hierarchical model summarizing the cellular and molecular differences in P7 *Fmr1* KO mice. This model integrates multi-level findings from lipidomic, transcriptomic, and cellular analyses in the P7 *Fmr1* KO mouse brain. At the base of the hierarchy, reductions in specific lipid species (PC 38:4 and PC 38:6) are observed in both brain and serum. Due to the limited strength of this experiment, this step in the model remains tentative and is denoted with a question mark. These deficits are associated with increased astrocyte-to-microglia communication via the EphA and SEMA pathways. At the cellular level, transcriptomic profiling reveals increased synaptic engulfment by glia, as well as decreased synaptic translation and microtubule stability in neurons. These molecular and cellular perturbations are accompanied by structural changes in synaptic architecture, including reduced RGC bouton size and decreased post-synaptic puncta size. Together, these results point to lipid dysregulation as a central mechanism underlying widespread cellular and synaptic abnormalities in *Fmr1* KO mice.

The results of this study expand the understanding of how FMRP loss impacts brain development by revealing a coordinated shift in glial and neuronal populations that supports aberrant synaptic pruning. While FMRP has been well studied for its role in regulating mRNA translation at the synapse, this study demonstrates that the absence of FMRP leads to defects in synaptic engulfment and circuit refinement. In parallel, transcriptomic, and cell communication analyses implicate disturbed lipid signaling pathways between astrocytes and microglia, with EphA and SEMA pathways emerging as key axes of interaction. Initial lipidomic analyses suggest a possible disruption of lipid-associated signaling,^66^ raising the hypothesis that changes in membrane composition could influence receptor organization. However, given the limitations of the lipidomic data, this mechanism is best viewed as preliminary. Together, these data support a model in which transcriptional and intercellular signaling changes drive synaptic dysfunction in FXS, with lipid dysregulation as a potential contributing factor.

The increase of EphA-associated lipid species through Lovastatin treatment suggests that the molecular disruptions associated with FMRP loss may be modifiable. However, given the exploratory nature of this analysis, the absence of a vehicle control, and the modest magnitude of the effects, these results do not establish a direct restoration of lipid raft structure or EphA receptor function. Drug intervention may also correct abnormalities in astrocyte-microglia communication and synaptic development, but future studies will be necessary to determine whether such lipid restoration is sufficient to rescue functional outcomes.

Despite these promising findings, several limitations should be acknowledged. This study focused on a single developmental time point, postnatal day 7, limiting insight into how the observed molecular and cellular changes progress over time or contribute to long-term neurodevelopmental outcomes. In addition, the functional and behavioral relevance of these molecular changes remains unclear, as this investigation did not include assessments of cognitive, social or sensory phenotypes. Addressing these gaps in future studies, particularly through longitudinal analyses, receptor-specific assays, and behavioral evaluations, will be critical for determining the relevance of glial-mediated mechanisms in FXS.

## AUTHOR CONTRIBUTIONS

ML, MSH, and CAM conceived the mouse experiments. Mouse eye experiments were performed and analyzed by ML (husbandry; synapse labeling and imaging; RGC segregation assay; lipidomics), with assistance from AV and MW. Single-nucleus RNA sequencing experiments were performed by ML and LC, with support from MG. Single-nucleus RNA seq data processing, clustering, pseudobulk and differential expression analysis, Gene Ontology Analysis, Cell-Cell communication analysis and total interaction potential (TIP) calculation were performed by LS, AY, FP, and VM. LS and ML designed the figures and wrote the manuscript; LS, ML, AY, FP, MSH, VM, and CAM made intellectual contributions and edited the manuscript.

## ACKNOWLEDGEMENTS

We thank all members of the Mason, Greenburg, Shirasu-Hiza, and Menon labs for support, discussions, and feedback. Funding for this research came from the NIH T32GM141882 (LS); T32EY013933 (ML), R35GM127049 (MSH); R01EYR01EY012736 (CAM); R21NS122366 (MSH, CAM). Simons Foundation Autism Research Initiative (SFARI) Explorer Award (CM, MSH, ML) and an Antonio Champalimaud Vision Award (CM).

