## Supplemental FIgures for "Disrupted glial-mediated synaptic refinement in Fragile X syndrome"

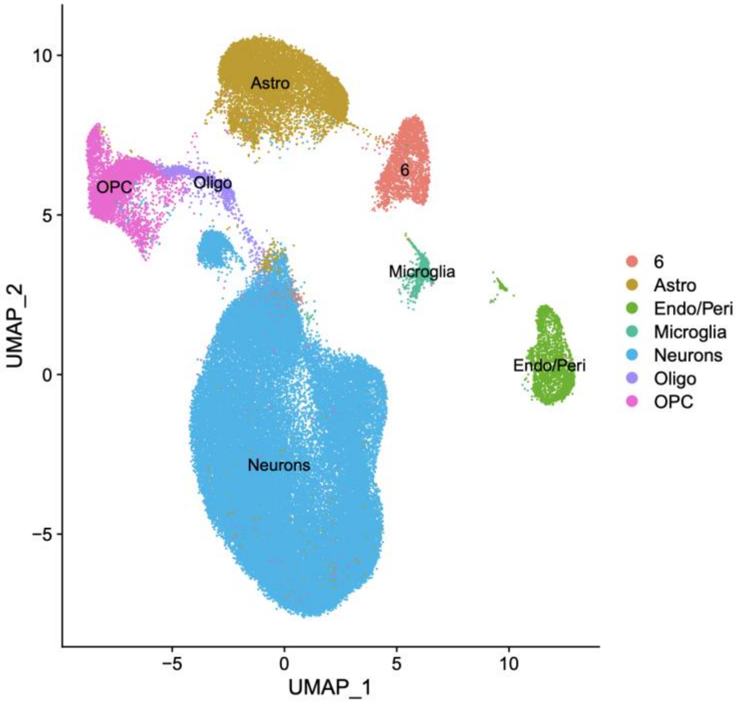


**Figure S1. Cell-type clustering of snRNAseq profiles from WT and *Fmr1* KO dLGN.** Uniform Manifold Approximation and Projection (UMAP) visualization of single-nucleus RNA sequencing data from dorsal lateral geniculate nucleus (dLGN) tissue pooled across WT and Fmr1 KO mice at P7. Unsupervised clustering revealed discrete transcriptionally defined cell populations corresponding to major neural and glial cell types, including neurons, astrocytes, oligodendrocytes (Oligo), oligodendrocyte precursor cells (OPC), microglia, and endothelial/pericyte populations (Endo/Peri). Cell identities were assigned based on canonical marker gene expression. The clear separation of clusters indicates robust transcriptional distinction across cell types and supports downstream pseudobulk, differential expression, and pathway analyses.

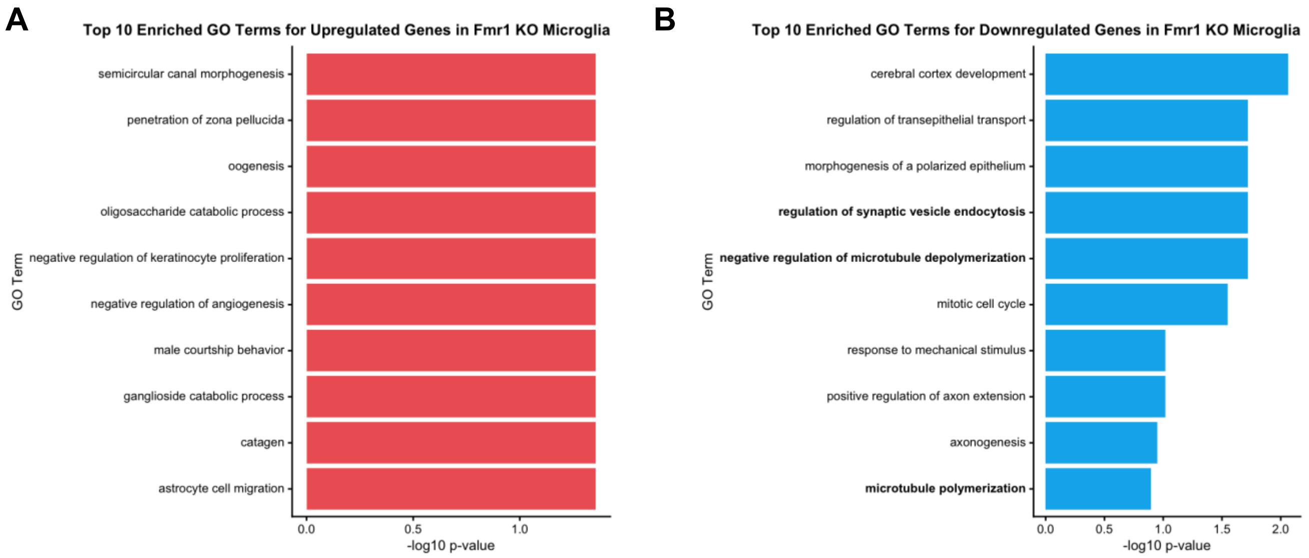


**Figure S2: Gene ontology (GO) analysis of *Fmr1* KO microglia genes. (A)** Top 10 GO terms enriched in upregulated microglial genes, showing increased expression of pathways related to astrocyte cell migration, consistent with a more reactive phenotype. GO enrichment was performed using the TopGO R package with the classic Fisher algorithm and a significance threshold of $p < 0.05$ **(B)** Top 10 GO terms among downregulated genes in *Fmr1* KO microglia, showing enriched regulation of synaptic vesicle endocytosis, negative regulation of microtubule depolymerization, and microtubule polymerization, indicative of dysregulation of cytoskeletal and trafficking pathways that support synaptic engulfment and refinement.

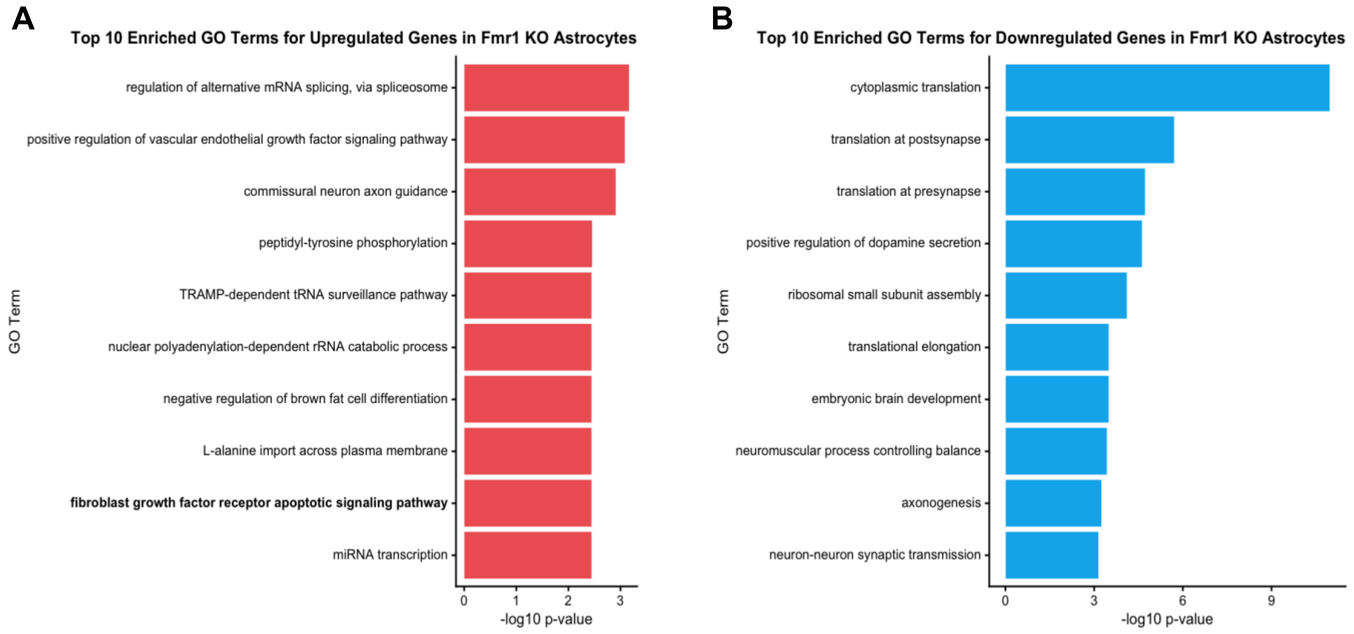


**Figure S3: Gene ontology (GO) analysis of *Fmr1* KO astrocyte genes. (A)** Top 10 GO terms enriched among genes upregulated in *Fmr1* KO astrocytes, including pathways related to RNA processing, vascular growth factor signaling, and axon guidance. These categories suggest altered regulation of transcriptional and signaling programs in astrocytes lacking FMRP. GO enrichment was performed using the TopGO R package with the classic Fisher algorithm and a significance threshold of $p < 0.05$. **(B)** Top 10 GO terms enriched among genes downregulated in *Fmr1* KO astrocytes, showing strong enrichment for cytoplasmic and synaptic translation, ribosomal assembly, and neurodevelopmental processes. These findings indicate suppression of pathways that support protein synthesis and synaptic function, consistent with impaired astrocytic contributions to synaptic maturation and neuronal support.

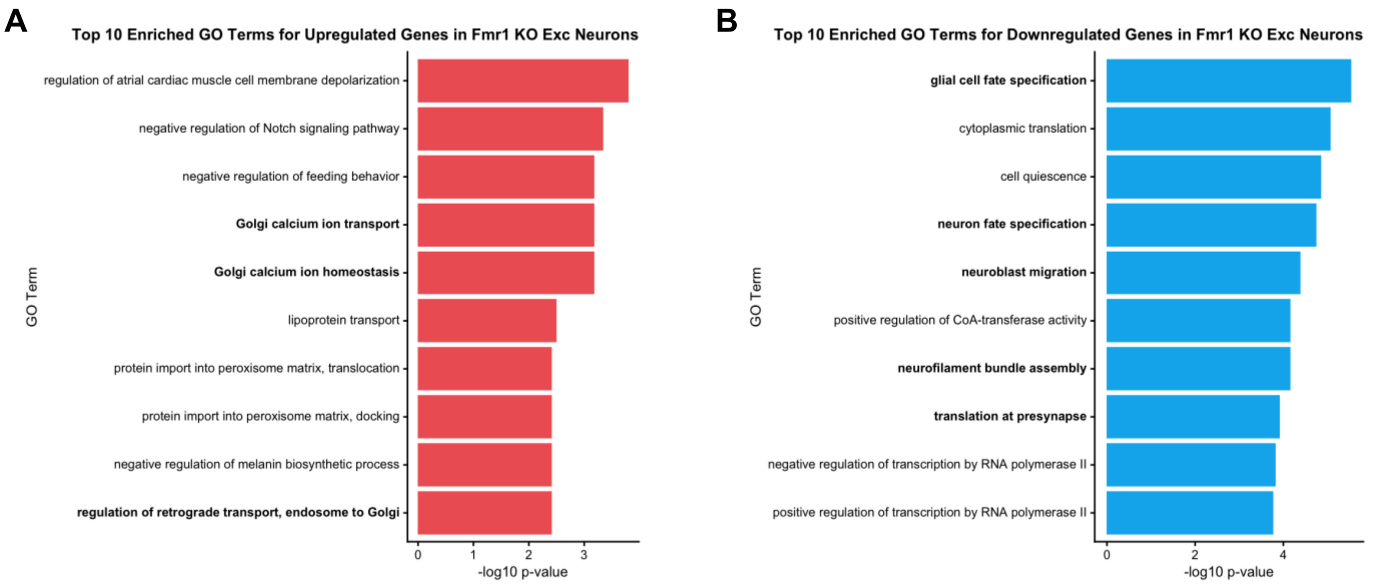


**Figure S4: Gene ontology (GO) analysis of *Fmr1* KO excitatory neuron genes. (A)** Top 10 GO terms enriched among genes upregulated in *Fmr1* KO excitatory neurons. Enriched categories include intracellular transport pathways and Golgi-associated calcium regulation, suggesting altered vesicle trafficking and membrane signaling processes in the absence of FMRP. GO enrichment was performed using the TopGO R package with the classic Fisher algorithm and a significance threshold of $p < 0.05$. **(B)** Top 10 GO terms enriched among genes downregulated in Fmr1 KO excitatory neurons, including glial and neuronal cell fate specification, neuroblast migration, synaptic translation, and neurofilament bundle assembly. These findings indicate suppression of pathways involved in neuronal maturation, cytoskeletal organization, and synapse-associated protein synthesis, consistent with impaired synaptic development in *Fmr1* KO mice.

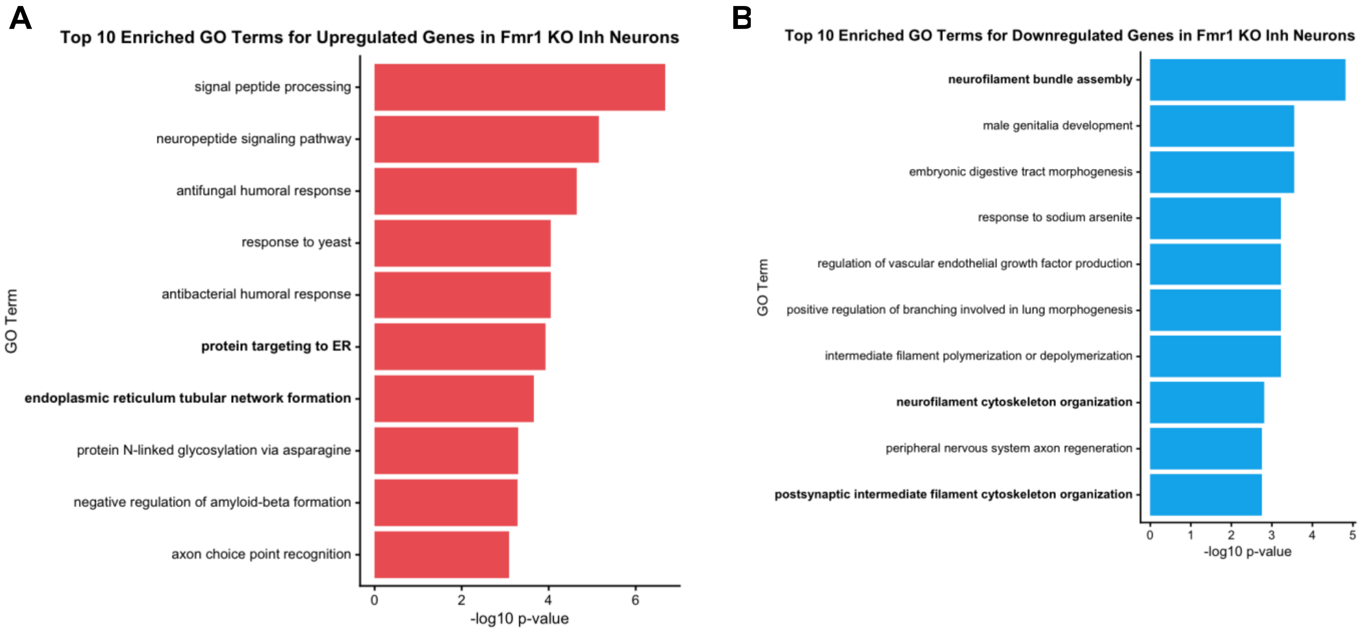


**Figure S5. Gene ontology (GO) analysis of *Fmr1* KO inhibitory neuron genes. (A)** Top 10 GO terms enriched among genes upregulated in Fmr1 KO inhibitory neurons. Enriched pathways include signal peptide processing, neuropeptide signaling, immune-related responses, and endoplasmic reticulum organization, suggesting altered peptide processing and intracellular protein trafficking in the absence of FMRP. GO enrichment was performed using the TopGO R package with the classic Fisher algorithm and a significance threshold of $p < 0.05$. **(B)** Top 10 GO terms enriched among genes downregulated in Fmr1 KO inhibitory neurons, including neurofilament bundle assembly, neurofilament cytoskeletal organization, and peripheral nervous system axon regeneration. These findings indicate suppression of pathways supporting cytoskeletal stability and axonal structure, consistent with altered neuronal maturation and structural support in Fmr1 KO inhibitory neurons.

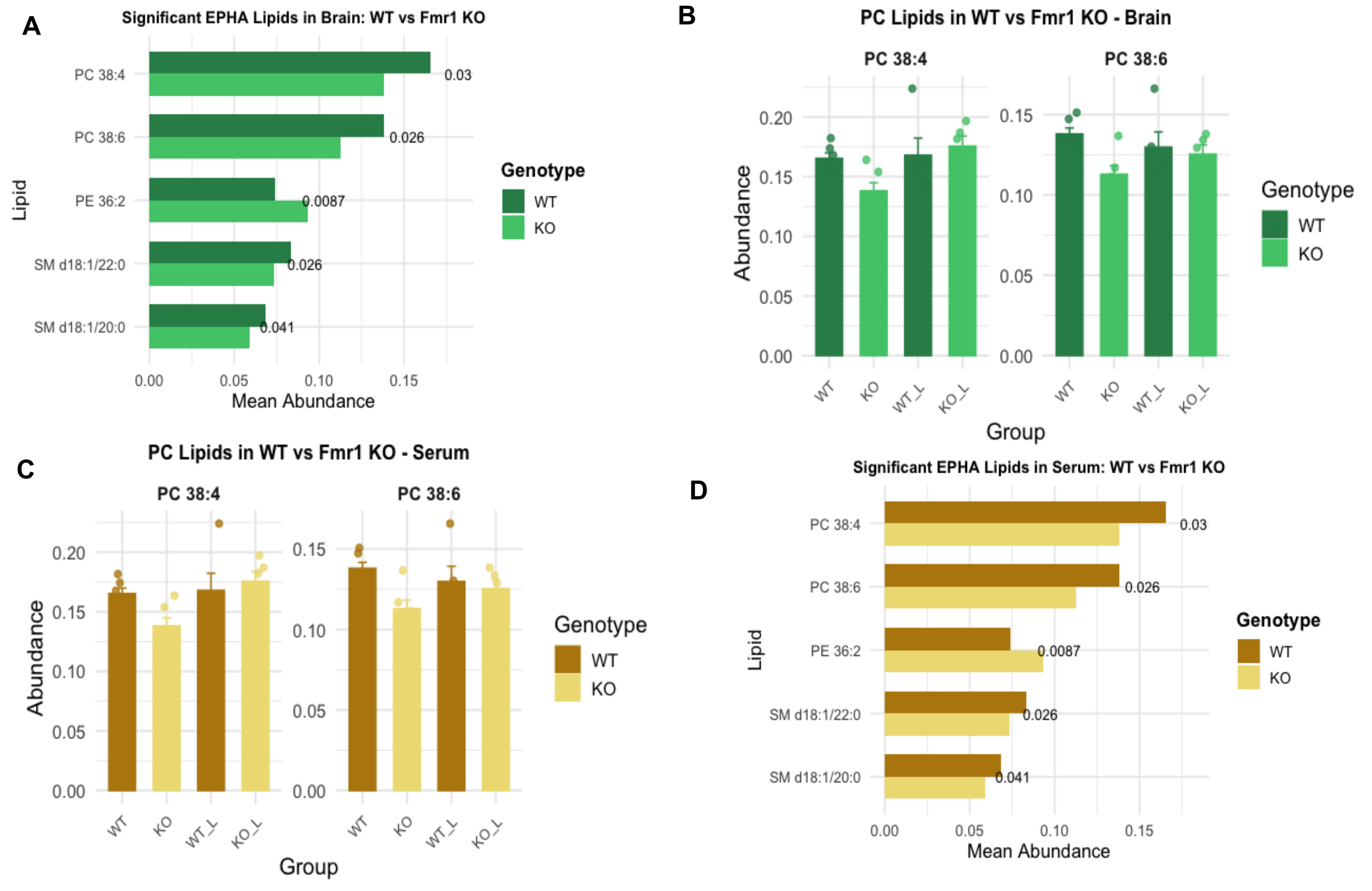


**Figure S6. EPHA-associated lipid deficits in *Fmr1* KO mice are reversed by Lovastatin treatment. (A, C)** Lipidomic profiling of brain **(A)** and serum **(C)** at P7 revealed significantly reduced levels of several EPHA-associated lipid species in *Fmr1* KO mice compared to WT controls, including phosphatidylcholines (PC 38:4, PC 38:6), phosphatidylethanolamine (PE 36:2), and sphingomyelins (SM d18:1/22:0, SM d18:1/20:0). These lipids are enriched in membrane microdomains such as lipid rafts, which regulate EPHA receptor localization and signaling. **(B, D)** Daily intraperitoneal injection of Lovastatin from P4–P6 restored the abundance of key EPHA-associated lipids—PC 38:4 and PC 38:6—in both brain **(B)** and serum **(D)** of *Fmr1* KO mice by P7. Data in **(A)** and **(C)** are represented as mean ± SEM and were analyzed using unpaired two-tailed Wilcoxon tests. Data in **(B)** and **(D)** are shown as mean ± SEM; individual points represent biological replicates.
